# Short 2′-O-methyl/LNA oligomers as highly-selective inhibitors of miRNA production *in vitro* and *in vivo*

**DOI:** 10.1101/2023.06.28.546821

**Authors:** Eloina Corradi, Natalia Koralewska, Marek C. Milewski, Linda Masante, Ryszard Kierzek, Marek Figlerowicz, Marie-Laure Baudet, Anna Kurzynska-Kokorniak

## Abstract

MicroRNAs (miRNAs) that share identical or near-identical sequences constitute miRNA families and are predicted to act redundantly. Yet recent evidence suggests that members of the same miRNA family with high sequence similarity might have different roles and that this functional divergence might be rooted in their precursors’ sequence. Current knock-down strategies such as antisense oligonucleotides (ASOs) or miRNA sponges cannot distinguish between identical or near identical miRNAs originating from different precursors to allow exploring unique functions of these miRNAs. We now develop a method based on short 2′-OMe/LNA-modified oligonucleotides to selectively target specific precursor molecules and ablate the production of individual members of miRNA families *in vitro* and *in vivo*. Using the highly conserved *Xenopus* miR-181a family as a proof-of-concept, we demonstrate that 2′-OMe/LNA-ASOs targeting pre-miRNA apical region elicit a precursor-selective inhibition of mature miRNA-5p production. The levels of miRNAs released from the 3′-arm of these precursors are not reduced, suggesting that our approach is also arm-selective. Overall, we show that this strategy can be successfully applied *in vivo* to achieve high target selectivity to study identical or highly similar miRNAs stemming from different precursors.

## INTRODUCTION

MicroRNAs (miRNAs) represent an evolutionarily conserved class of small non-coding RNAs that function in post-transcriptional gene silencing by base-pairing to the cognate messenger RNAs (mRNAs) [1]. miRNAs are important regulators of various physiological processes, including animal development [2–6]. They are also involved in pathogen-response and viral infection [7–9]. miRNA expression levels change under multiple physiological and pathological conditions [10–12], making them potential biomarkers and druggable targets [13, 14]. Therefore, the ability to modulate miRNAs expression is important for uncovering their physiological roles through functional studies as well as for developing innovative therapeutic strategies.

The biogenesis of miRNA is a multistep process typically involving sequential cleavage of miRNA precursor molecules – pri-miRNA and pre-miRNA – by two RNase III enzymes, Drosha and Dicer, respectively [15]. First, in the nucleus, primary transcripts of miRNAs (pri-miRNAs) are processed by Drosha. This cleavage yields ∼60-nucleotide (nt) pre-miRNAs, which are exported into the cytoplasm, where they are further cut by Dicer endoribonuclease. Dicer cleaves both arms of the pre-miRNA hairpin to yield ∼22 base pairs-long miRNA-5p/miRNA-3p duplex. Next, one of the miRNA strands is incorporated into an RNA-induced silencing complex (RISC) where it serves as a guide molecule directing RISC to specific complementary gene transcripts to ultimately induce silencing of the target [16]. The strand discarded during RISC assembly (a passenger strand, miRNA*) is thought to undergo rapid degradation, although abundant accumulation of certain miRNAs* and their incorporation into RISC have been also reported [17, 18].

Experimental evidence shows that target specificity is primarily determined by nucleotides at position 2-7 of miRNA, called the “seed” sequence [19], and miRNAs with the same seed sequence are grouped in a “miRNA family”. Because of the shared seed, members of the same miRNA family are predicted to act redundantly and regulate a similar set of transcripts (the “seed” hypothesis). However, over the years this assumption has been challenged by several reports showing that individual family members can acquire specific functions [20–29]. In particular, miRNAs that have in common their 5′ sequences but differ in their 3′ ends, e.g. members of let-7 family, have been found to target largely distinct sets of sites within transcriptome [23–25]. This underscores the contribution of the region outside the seed to miRNA function. Remarkably, for some miRNA families, the functional specificity of individual miRNAs seems to depend not on their sequence *per se* but on the pri- and/or pre-miRNA apical loop sequence [26–28]. For instance, murine miR-181a-1-5p and miR-181c-5p differ by only one nucleotide (position 11th of the mature miRNA sequence) [26], and thus according to the “seed” hypothesis, they should have similar roles in cells. However, it was demonstrated that only miR-181a-1-5p and not miR-181c-5p promotes T cell development when ectopically expressed in thymic progenitor cells [26]. Through mutagenesis and pre-miRNA-181a-1/pre-miRNA-181c regions swapping experiments, it was further shown that the difference between miR-181a-1 and miR-181c activities is associated with pre-miRNA loop sequences and not mature miRNA sequences [26]. The exact mechanism of this phenomenon is not fully understood. A plausible explanation is that regulatory proteins binding the pri-miRNA/pre-miRNA apical loop are affecting the generation of the mature miRNA, similarly to Lin28 binding pre-let-7 loop and inhibiting its maturation [30–32] or TDP-43 which on the contrary promotes miRNAs biogenesis by binding pri- and pre-miRNA terminal loop [33]. Alternatively, it is possible to hypothesize an apical loop-dependent mechanism where the loop structure itself promotes a differential sorting of miRNAs into RISCs. This latter hypothesis is supported by studies showing that pre-miRNAs structural features within the hairpin stem (i.e. internal loop or bulges) are responsible for the selective sorting of miRNAs to functionally distinct RISC complexes [34, 35].

The importance of the pre-miRNA sequence for mature miRNA function is also illustrated by the differential distributions of precursors encoding identical 5p form. For example, while both pre-miR-138-1 and pre-miR-138-2 can give rise to miR-138-5p, in rodent oligodendrocytes this miRNA originates predominantly from pre-miR-138-1, whereas in neurons – from pre-miR-138-2 [36]. In addition, in the African clawed frog *Xenopus laevis*, pre-miR-181a-1 is significantly more abundant than pre-miR-181a-2 in developing axons, yet both give rise to miR-181a-5p [5]. As in *Xenopus*, human pre-miR-181a-1 and pre-miR-181a-2 are the sources of identical miR-181a-5p. They are also co-transcribed with pre-miR-181b-1 and pre-miR-181b-2 respectively, which give rise to identical miR-181b-5p. Interestingly, in human natural killer (NK) cells, the levels of pre-miR-181a-1/b-1 vs pre-miR-181a-2/b-2 are differentially regulated during NK lineage development and in response to various cytokines [37]. It cannot be ruled out that in the abovementioned cases, differential expression of the pre-miRNAs may serve the regulation of functionally different miRNAs-3p. Nevertheless, these differing miRNA-3p forms might still represent also a way to regulate the selection of corresponding miRNAs-5p, i.e. by impacting their loading in the RISC complex [34, 35]. Taken together, these data suggest a pivotal role for precursor identity and processing in regulating mature miRNA fate. Yet the functional significance of this intriguing phenomenon remains unknown.

Unfortunately, no critical insight has been gained beyond these initial studies because to date there were no tools to study the function of identical or near-identical miRNAs stemming from different precursors. The most common approach to investigate miRNA functions involves the loss-of-function (LOF) methods, including: (i) genetic knockout of a specific miRNA; (ii) miRNA sponges [38]; (iii) antisense oligonucleotide inhibitors (ASOs) [39–41]. Of these methods, genetic knockout and miRNA sponges are not suitable to tease apart precursors generating identical or near-identical miRNAs. Knockout models in which only specific miRNA genes are disrupted are challenging to generate without perturbation of other genes within a miRNA cluster. miRNA sponges, which are exogenously introduced transcripts that contain multiple binding sites complementary to the seed region of a miRNA of interest [42], are commonly designed to inhibit the activity of an entire family of miRNAs sharing a common seed. Thus, they are not a good choice for the LOF studies of precursors carrying identical or near identical miRNA sequences [43]. The third method, ASOs, gives a unique opportunity to target specific pre-miRNA regions. ASOs can be also delivered to specific cells at the organismal level, e.g. by microinjection or electroporation [44], opening the possibility to study cell- or tissue-specific roles of miRNA. We previously used morpholino (MO) ASOs in the *X. laevis* RGC axon model to target miRNA precursors, yet we designed them to indiscriminately block both pre-miR-181a-1 and pre-miR-181a-2 processing [5] exploiting their ability to be broad inhibitors. Overall, no ASO strategy has been hitherto specifically devised to address the question of precursor contribution to miRNA function.

We, therefore, sought to design a new ASO-based strategy to target miRNA precursors. We optimized ASO targeting efficiency affinity, specificity, and stability by shortening ASO length to 13-nt and integrating 2′-O, 4′-C-methylene bridge (LNA) [40] and 2′-O-methyl (2′-OMe) modifications [39]. As a proof-of-principle, we tested our strategy, both *in vitro* and *in vivo*, by targeting precursors of miR-181a-5p in *Xenopus laevis*, one of the most widely-used non-mammalian systems to study fundamental biological processes *in vivo* [45–47], including miRNAs biogenesis and functions [5, 48–51]. miR-181a-5p is a member of the highly conserved vertebrate miR-181a family [52] and it originates from two precursors: pre-miR-181a-1 and pre-miR-181a-2 which differ in the size and sequence of their apical loops and give rise to distinct miRNAs-3p (miR-181a-1-3p and miR-181a-2-3p, respectively). miR-181a-5p is the second most abundant miRNA in retinal ganglion cells (RGC) axons of *X. laevis* [49], and has an important role in RGC axon development, and in the formation of a fully functional visual system [5]. Although miR-181a-1-3p and miR-181a-2-3p are also abundant in RGC axons (i.e. the 30^th^ and the 9^th^ most abundant axonal miRNAs respectively) [47], it remains unclear whether they have a function in these processes.

We demonstrate here that 13-nt 2’-OMe/LNA ASOs achieve the efficient inhibition of miR-181a-5p production in a precursor selective manner, discriminating between the two highly similar pre-miR-181a-1 and pre-miR-181a-2. This strategy does not reduce the production of miRNAs from the opposite pre-miRNA arm (i.e. miRs-181a-1-3p and miRs-181a-2-3p). In addition, we show that these short ASOs can be used *in vivo* without any cytotoxic effect and with higher precursor- and arm-selectivity than longer 24-25-nt MOs. Our results indicate that short 13 nt 2’-OMe/LNA-modified oligonucleotides designed to target the variable apical region of pre-miRNA spanning Dicer cleave site and the loop region can be successfully used for highly selective inhibition of miRNA production at the level of both the individual pre-miRNA (arm-selective inhibition) and miRNA family (precursor-selective inhibition).

To the best of our knowledge, this is the first study exploring the use of such short 2′-OMe/LNA ASOs to selectively inhibit the release of identical miRNAs from two paralogous precursors. The proposed approach offers a potent tool to significantly enhance the investigation of the functional diversification within miRNA families and study the contribution to biological processes and disease of individual miRNAs stemming from different precursors.

## RESULTS AND DISCUSSION

### Design of short 2′-OMe/LNA oligomers targeting miR-181a-5p precursors

#### Design rationale

The design of potent and specific ASOs requires in-depth consideration of oligonucleotide’s parameters such as: (i) target affinity, i.e. the strength by which the ASO and the target bind, (ii) target specificity, i.e. how restrictive the ASO is in its choice of the target, and (iii) ASO stability in a cell. It is essential to balance target affinity against the risk of mismatched off-target hybridization that both comes with increasing the oligomer length. The known ASO-based strategies targeting pre-miRNAs typically involve ∼20 nt oligonucleotides [53], but this design has significant shortcomings. In particular, such long oligomers have a higher potential to adopt secondary or tertiary structures that leave them unable to hybridize to the target, reducing their efficiency. In addition, they are more likely to generate non-specific effects by mismatched hybridization, i.e. partial binding to unintended targets. It is estimated that in a typical higher eukaryotic mRNA pool of about 10^4^ different mRNA species of an average length of 2 x 10^3^ bases, the shortest sequence that is likely to be unique is 13 nt long [54].

As a model for developing our ASO-based approach, we used two *X. laevis* precursors: pre-miR-181a-1 (62 nt) and pre-miR-181a-2 (67 nt). pre-miR-181a-1 and pre-miR-181a-2 differ in the size (11 nt vs 16 nt) and sequence of their apical loops, and both give rise to the identical miR-181a-5p, and to distinct miRNAs-3p: miR-181a-1-3p and miR-181a-2-3p, respectively (Supplementary Figure S1A). We aimed to inhibit the production of miR-181-5p from a specific precursor, without perturbing the production of the corresponding miRNA-3p. Taking into account the facts that the shortest sequence predicted to occur only once in the vertebrate mRNA pool is 13-nt long [54] and that the apical loop region is the most divergent part of both precursors, with only 67% sequence identity and 5-nt length difference between pre-miR-181a-1 and pre-miR-181a-2 (Supplementary Figure S1A), we designed a pair of oligomers to separately target each precursor: (i) 13-nt ASO overlapping Dicer 5′ cleavage site and partially spanning the apical loop, and (ii) ASO complementary to the entire apical loop of the precursor.

#### In silico analysis

To choose the optimal sequence for the precursor-specific ASOs, we used an in-house pipeline created to design oligomers of defined length that also display the best hybridization parameters, including minimal self- and cross-hybridization (Figure 1A). Both pre-miR-181a-1 and pre-miR-181a-2 adopt hairpin structures with very similar stems (92% identity i.e. 47 out of 51 nt) but with very different size and nucleotide composition of apical loops (Figure 1B, Supplementary Figure S1). Pre-miR-181a-1 has an 11-nt loop with uniform nucleotide content: 45.5% G/C and 54.5% A/U, whereas pre-miR-181a-2 has a larger, 16-nt loop, with only 19% of G/C and 81% of A/U (Supplementary Figure S1A). Thus, ASOs complementary to the entire apical loop of these pre-miRNAs differ in length. Altogether, we designed four ASOs, which we named AL1-4 (Figure 1C). AL1 and AL2 (both 13 nt) were complementary to pre-miR-181a-1: AL1 overlapped Dicer 5′ cleavage site, and AL2 was complementary to the entire hairpin loop (Figure 1C, Supplementary Figure S1A-B). AL3 (13 nt) and AL4 (18 nt) were complementary to pre-miR-181a-2: AL3 overlapped Dicer 5′ cleavage site, whereas AL4 (18 nt) was complementary to the entire hairpin loop of pre-miR-181a-2 (Figure 1C, Supplementary Figure S1A-B). To compensate for the relatively short length of the oligomers, we combined the benefits of 2′-O, 4′-C-methylene bridge (LNA) [40] and 2′-O-methyl (2′-OMe) modifications [39]. Because of their remarkable hybridization properties, the introduction of LNA and 2′-OMe modifications allows ASO length to be shortened while preserving target affinity and selectivity. For example, it has been reported that LNA oligomers even as short as 8 nt can achieve sufficient binding affinity to efficiently regulate target miRNA [55]. Moreover, LNA has been proven to maximize mismatch discrimination by destabilizing non-fully complementary duplexes composed of ASOs and unintended targets [56, 57]. Likewise, 2’-OMe oligonucleotides have been also shown to bind RNA targets with much higher affinity than corresponding unmodified oligonucleotides [58, 59], with the maximal beneficial effect observed for targets of 16 nt or less [58]. Additionally, 2’-OMe/LNA modifications improve the nuclease resistance and cellular half-life time of ASOs [59, 60]. Therefore, each AL oligomer consisted of 2′-OMe/LNA nucleotides, with LNA positioned near the ends and in the middle of the strand (Figure 1C). Such a location of the LNA modifications is known to increase oligomers’ affinity to the target RNA [61, 62]. One of the parameters most often used when comparing the hybridization affinity of the oligonucleotides is the change of the free Gibbs energy (ΔG) [63]. The lower (more negative) the value, the stronger the interaction between the oligonucleotides. Using EvOligo [64] we calculated ΔGs for the duplexes formed by AL-ASOs and their respective miR-181a precursors, as well as AL-ASOs and other pre-miRNAs (potential off-targets identified within known *X. laevis* precursors). We found that ΔG for AL:pre-miR-181a-1/-2 duplexes were all below -21 kcal/mol (Figure 1C, Supplementary Table S1), which was at least 1.5 times lower than for AL-ASOs pairs with any other pre-miRNA. These results suggested stronger on-target than off-target hybridization affinity of the designed AL-ASOs (Supplementary Table S1).

**Figure 1.**
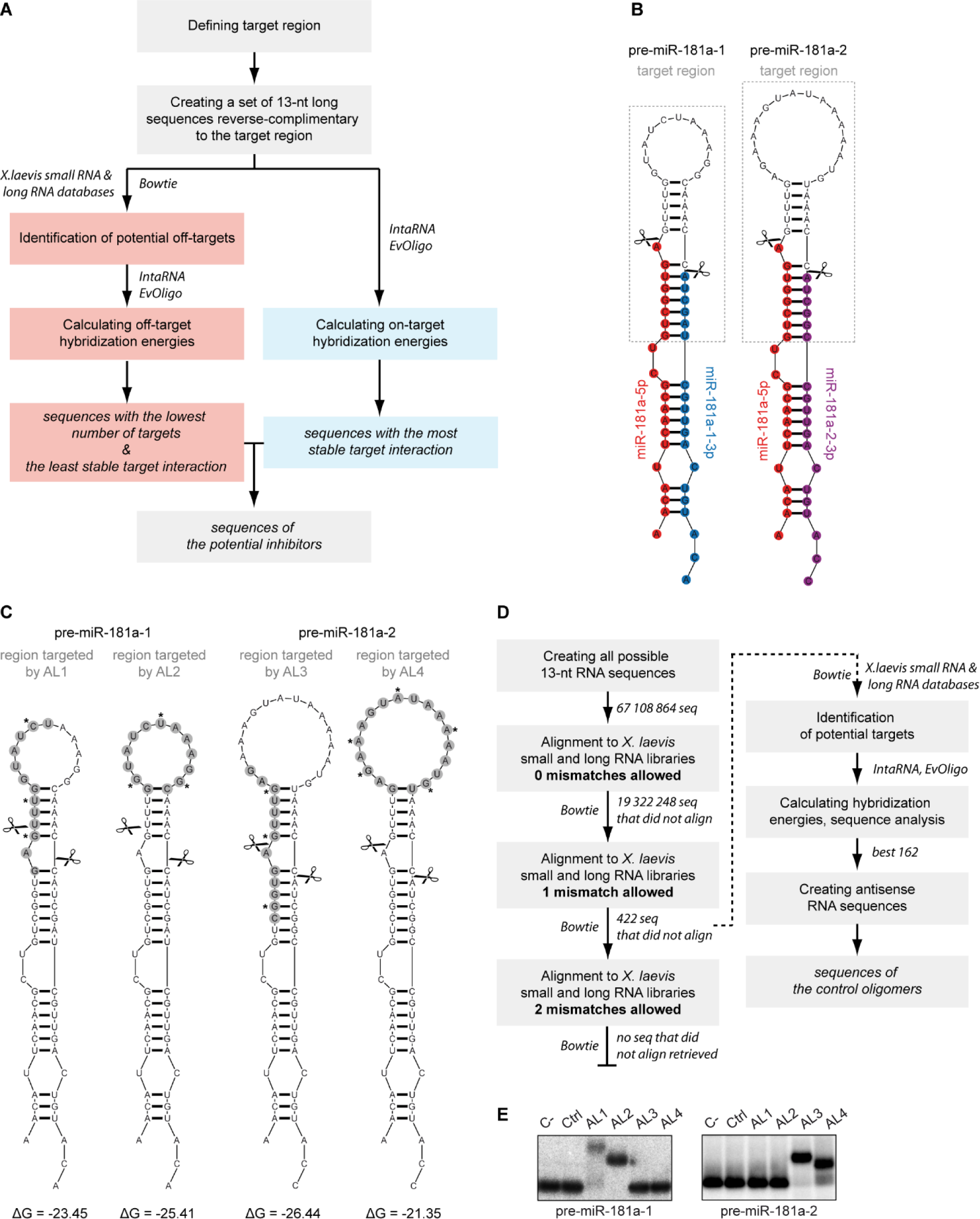
Designing ASOs targeting the apical region of pre-miRNA. **(A)** Schematic representation of the inhibitors design workflow. **(B, C)** Predicted secondary structures of pre-miR-181a-1 (*left*) and pre-miR-181a-2 (*right*). Scissors indicate Dicer cleavage sites. (B) Mature miRNA sequences are coloured and grey boxes indicate the target sequence used in the ASOs design process. (C) Grey circles mark regions targeted by the selected oligomers (AL1-4), asterisks indicate the position of LNA within the AL-ASO sequence, free Gibbs energy change, ΔG [kcal/mol] of AL-ASO:pre-miRNA complexes is given. **(D)** Schematic representation of the control oligomers design workflow. **(E)** Native gel analysis of the interaction between 5′-^32^P-labeled pre-miRNA and tested oligomers (Ctr1, AL1-4). Abbreviations: AL, apical loop; Ctrl, control oligomer; C-; pre-miRNA with no other oligomer added.

Additionally, we designed a negative 13-nt control oligomer (Figure 1D). Typically, control oligomers used in ASO-based experiments possess either a scrambled sequence or a sequence generated randomly. To minimize the risk of the off-target effects resulting from unintended hybridization of the control oligomers to endogenous RNAs, we took a different approach and looked for 13-nt sequences that display the least complementarity to any known *X. laevis* transcripts (Figure 1D). To this end, first we generated a set of all possible 13-nt sequences randomly obtained by the combination of the 4 nucleotides (a total of 4^13^ sequences). Second, we removed from this set all sequences identical with any known *X. laevis* transcript (∼70% of possible 13-nt sequences were excluded). Third, to further strengthen the selection conditions, we excluded all the sequences that were still aligned to *X. laevis* transcripts when one mismatch was allowed (i.e. 12 out of 13 nt were identical), which resulted in narrowing the set of analyzed sequences that did not have a perfect match to 422. When two mismatches were allowed (i.e. 11 out of 13 nt were identical), all the 422 sequences had a match within *X. laevis* transcripts (Figure 1D). This showed that no further strengthening of the selection conditions was possible. Next, we filtered the set of 422 sequences, removing those with characteristics unfavorable for ASOs, i.e. three (or more) consecutive identical nucleotides or nucleotide pairs, or the tendency to form homo-duplex. After this step, we obtained a pool of 162 sequences for which the antisense sequences were created. These antisense sequences represented the set of potential negative control oligomers for *X. laevis*, as they do not have any known fully complementary or single-nucleotide mismatch targets in the *X. laevis* transcriptome (Supplementary Table S2). From this set of equally-well scored sequences, we arbitrarily chose one sequence to obtain the control oligomer (named “Ctrl”) used for further experiments (Supplementary Table S2).

#### In vitro validation of AL1-4 target selectivity

ASOs exert their function by binding to the complementary sequence within the target nucleic acids. Thus, when selecting ASO-based inhibitors for *in vivo* studies, it is useful to first assess the efficiency and specificity of their binding to the target RNA/DNA *in vitro*. Here, to study AL:pre-miRNA interactions, we used gel electrophoresis under non-denaturing conditions (native gel electrophoresis). The band shift revealed that AL1 and AL2 formed stable duplexes with pre-miR-181a-1 but not with pre-miR-181a-2, whereas AL3 and AL4 bound to pre-miR-181a-2 but not to pre-miR-181a-1 (Figure 1E). Ctrl did not form complexes with either of the precursors. Densitometry analysis indicated that AL2 bound 100% of pre-miR-181a-1, whereas AL1 bound only ∼75% of this precursor; AL3 bound ∼95% of pre-miR-181a-2, whereas AL4 bound only ∼70% of it. These results are in line with the predicted free energy values of the RNA:RNA complexes (Supplementary Table S1), which also pointed to the possible difference in target binding efficiency. The free energy values calculated for AL2:pre-miR-181a-1 complex (−25 kcal/mol) and the AL3:pre-miR-181a-2 complex (−26 kcal/mol) were 2-5 kcal/mol lower than the values of AL1:pre-miR-181a-1 (−23 kcal/mol) and AL4:pre-miR-181a-2 (−21 kcal/mol), respectively. It is also important to note that the target region for AL3 in pre-miR-181a-2 (Figure 1C) differs by only 2 nt from the corresponding fragment in pre-miR-181a-1 (Figure 1C). Since AL3 bound to pre-miR-181a-2 but not to pre-miR-181a-1, this indicates that it was able to distinguish between two highly similar targets. This suggests that ≥2-nt mismatch is likely sufficient to prevent a real off-target stable interaction in the case of 13-mer ASOs. This observation is also in favor of our control design strategy, since the Ctrl oligomer has potential targets in the *X. laevis* transcriptome but all of them have at least two mismatches (Figure 1D, Supplementary Table S2).

Taken together, these results confirmed that *in vitro* all of our designed oligomers displayed high mismatch/target discrimination and high specific hybridization affinity towards the desired pre-miRNAs.

### Assessment of the inhibitory potential of designed oligomers *in vitro*

#### Analysis of the efficiency and selectivity of AL1-4 in X. laevis cytosolic extracts

We next investigated the inhibitory efficiency and target selectivity of AL1-4 in the simple *X. laevis* model using cellular cytosolic extracts from embryos, which contain endogenous *X. laevis* Dicer (Figure 2A). In these studies, we used extracts from stage 44 tadpoles, where miR-181a-5p expression is the highest across development [65]. We reasoned that at this stage, the cytosolic concentration of the potential endogenous inhibitors targeting miR-181a precursors would be the lowest, which would better allow for unambiguous testing of ASO inhibitory potential.

**Figure 2.**
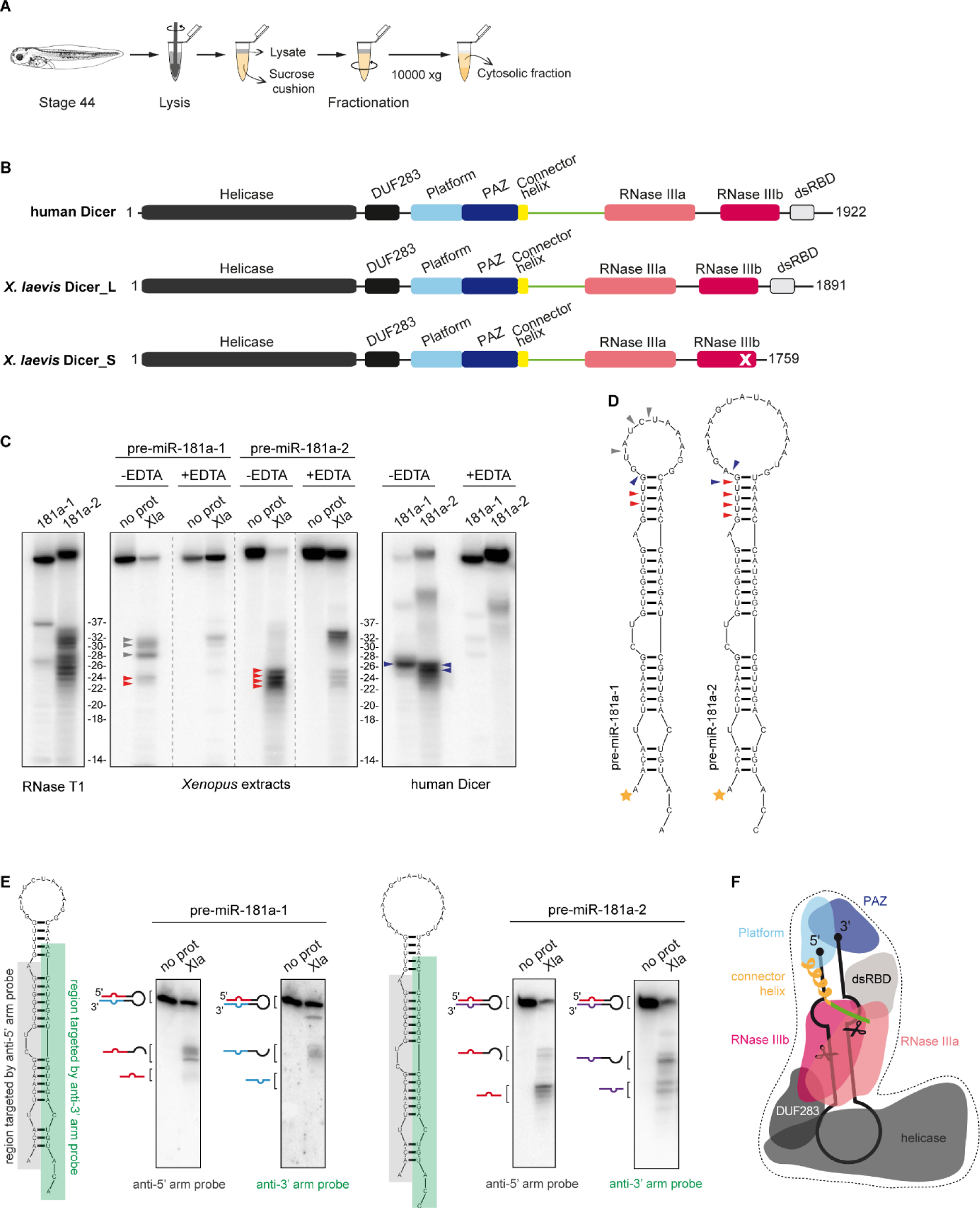
Characterization of the pre-miRNA cleavage pattern generated in cytosolic extracts. **(A)** Schematic of the experimental paradigm for cytosolic extracts from stage 44 *X. laevis* embryos. **(B)** Schematic representation of domain composition of human and *X. laevis* Dicer proteins. A cross-mark indicates amino-acid substitutions in the catalytic core of *X. laevis* Dicer S. **(C)** Representative gels showing 5′-^32^P-labeled pre-miRNA cleavage patterns upon incubation with *Xenopus* cytosolic extracts or human Dicer with or without EDTA. The ladder was created based on pre-miR-181a-1 and pre-miR-181a-2 cleavage patterns generated by RNase T1. Red and blue arrowheads indicate bands accepted as representing miR-181a-5p isoforms generated in *X laevis* extracts and by human Dicer, respectively. Gray arrowheads point to the bands marked as non-Dicer products. Reproducible results were obtained using different extract batches. **(D)** Schematic representation of the cleavage pattern of pre-miR-181a-1 and pre-miR-181a-2 established based on the PAGE results. Arrowheads as in (C); orange stars indicate 5′-^32^P-labeling. **(E)** Results of Northern blot analysis of the cleavage pattern of pre-miR-181a-1 and pre-miR-181a-2 in *X. laevis* extracts using probes targeting the 5′ or 3′ arm of the precursor. Pre-miRNA regions targeted by the probes are indicated and schematic representation of the pre-miRNA, AL-ASO and the cleavage products is given on left to the blots. N Abbreviations: Xla, stage 44 *X. laevis* cytosolic extracts; hDicer, human Dicer; no prot, no protein control; DUF283, domain of unknown function 283; PAZ, Piwi/Argonaute/Zwille domain; dsRBD, dsRNA binding domain. **(F)** Schematic representation of the tertiary structure of Dicer in a complex with pre-miRNA. Prepared based on cryo-EM structure obtained for human Dicer by Liu et al. [83]. Scissors indicate the sites of pre-miRNA cleavage by RNase III domains. Figure adapted from [84]. Color code as in (B). Abbreviations: DUF283, domain of unknown function 283; PAZ, Piwi/Argonaute/Zwille domain; dsRBD, dsRNA binding domain.

Allotetraploid *X. laevis* cells express two Dicer forms, named L and S, that are encoded on long and short chromosomes, respectively (Figure 2B, Supplementary Figure S2). Most Dicer enzymes are multidomain proteins consisting of a putative helicase domain, a domain of unknown function (DUF283), Platform, Piwi–Argonaute–Zwille (PAZ) domain, two RNase III domains (RNase IIIa and RNase IIIb), and a dsRNA-binding domain (dsRBD) [66]. *X. laevis* Dicer L is similar to Dicers of higher organisms, including humans, while Dicer S lacks the dsRBD domain and contains amino-acid substitutions within the catalytic center of the RNase IIIb domain, which should render it inactive (Figure 2B, Supplementary Figure S2) and presumably not involved directly in pre-miRNA processing. Dicer enzymes specifically hydrolyze phosphodiester bonds in double-stranded regions, the cleavage is Mg^2+^-dependent, and the products length range between ∼20-26 nt, depending on the species-specific, spatial orientation of the PAZ, Platform, and RNase III domains [66].

We first evaluated Dicer enzymatic activity in *X. laevis* cytosolic extracts using 5′-^32^P-labeled pre-miR-181a-1 and pre-miR-181a-2 as substrates, and recombinant human Dicer as a control. The ^32^P at the 5′-terminus of the substrate allowed us to track the release of miR-181a-5p. All reaction mixtures were analyzed by PAGE, and radiolabeled substrates and products were visualized by phosphor imaging (Figure 2C). Taking into consideration structural and biochemical determinants of RNA cleavage by Dicer, we considered RNA fragments as specific 5′ arm miRNA products (miRNA-5p) if they meet the following criteria: (i) their size was between 20 and 26 nt, (ii) their generation required cleavage within the stem of the precursor, and (iii) cleavage was abrogated or reduced in the presence of Mg2+-chelating agents (e.g. EDTA) (Figure 2C, D) [67]. To gain a more comprehensive look into the spectrum of the fragments generated in the reaction and visualize RNA beyond 5′-^32^P-labeled molecules, we conducted Northern blot assays using probes targeting either the 5′ or 3′ arm of the precursor (Figure 2E).

When examining the putative cleavage of both pre-miR-181a-1 and pre-miR-181a-2 by cytosolic extracts, we detected ∼22-26 nt products that diminished under the EDTA conditions (Figure 2C, D). We classified these fragments as miR-181a-5p isoforms (isomiRs). Such miRNA variants deviating from the canonical sequence by up to a few nucleotides are not uncommon for miR-181a-5p, as many as 39 isoforms of miR-181a-5p have been found in T-cell acute lymphoblastic leukemia samples [68]. For pre-miR-181a-1, we detected two miR-181a-5p isomiRs, 23 nt and 24 nt long, with the latter being more abundant (Figure 2C). For pre-miR-181a-2, we detected four variants of miR-181a-5p ranging from 22 to 25 nt, with 24-nt form being the most prevalent (Figure 2C). Densitometry analysis revealed that under the experimental conditions, miR-181a-5p (all isomiRs counted) were generated 4.6 times more efficiently from pre-miR-181a-2 than from pre-miR-181a-1 (Figure 2C, E, Supplementary Figure S3). Additionally, in the case of pre-miR-181a-1 we detected 28-32-nt fragments, which we termed pre-miRNA “halves” because of their length corresponded roughly to half of that of the precursor (Figure 2C, E). Gel densitometry showed that the pre-miRNA halves were produced 4 times more efficiently than miR-181a-5p isomiRs (Figure 2C, E).

When examining the putative cleavage of either precursor by recombinant human Dicer, we detected one dominant 26-nt form of miR-181a-5p generated in reactions involving pre-miR-181a-1, and two major isomiRs (25 and 26 nt) generated in the reactions with pre-miR-181a-2. For both precursors, we did not observe ∼28-30 nt fragments (pre-miRNA halves). This further pointed to the possibility that a cellular factor caused the appearance of pre-miRNA halves in *X. laevis* extracts rather than the intrinsic instability of pre-miR-181a-1 or Dicer activity. In general, isomiRs generated in *X. laevis* extracts were ∼2 nt shorter than those produced by the recombinant human Dicer (Figure 2C, D). The alignment of *X. laevis* and human Dicer amino acid sequences showed that the region connecting the PAZ and RNase III domains involved in “measuring” the size of the dicing product is 12 amino acids shorter in *X. laevis* enzymes as compared to the human one (Figure 2B, F, Supplementary Figure S2), which might explain the observed difference in the isomiRs size. It is also important to emphasize that the 2-nt difference between *X. laevis* and human Dicer products is consistent with the data collected in the miRBase, in which 21-nt *X. laevis* miR-181a-5p (MIMAT0046485) and 23-nt human miR-181a-5p (MIMAT0000256) have been reported.

Next, we used a similar approach to test the inhibitory potential of AL1-4, with each AL-ASO assayed at three concentrations: 0.1, 1, and 10 µM (Figure 3, Supplementary Figure S4). We first examined AL1 and AL2, which target pre-miR-181a-1. In reactions with AL1 but not AL2, the levels of miR-181a-5p were decreased and a longer, 36/37-nt product corresponding to the portion of the precursor left after miR-181a-3p excision was detected (Figure 3A, Supplementary Figure S4A). Densitometry analysis revealed that at the highest AL1 concentration (10 µM), the level of miR-181a-5p was reduced by ∼70% in comparison to the positive control reaction in which 5′-^32^P-pre-miR-181a-1 was incubated in extracts only (C+) or in reaction with Ctrl oligomer (Figure 3A). The inhibition of miR-181a-5p production was also dose-dependent (Figure 3A) with a significantly lower inhibition at 0.1 µM and 1 µM, compared to 10 µM (Figure 3A). Unexpectedly, in samples with AL2, the level of miR-181a-5p was greatly increased in comparison to the control reaction (Figure 3A, Supplementary Figure S4B). The accumulation of miRNA was positively correlated with the concentration of AL2: at the highest concentration of AL2 (10 µM), the relative level of miR-181a-5p was almost 3.5 times higher (Figure 3A, right panel) than in the positive control reactions (C+). Additionally, at the higher concentrations of AL2 (1 and 10 µM), we did not observe products identified as pre-miRNA halves. We conclude that AL2, which is complementary to the entire apical loop of pre-miR-181a-1 but does not overlap with Dicer cleavage sites (Figure 1C), may shield pre-miR-181a-1 from negative regulators acting on a loop region, and does not affect processing by Dicer. This scenario is further corroborated by the results from the control reactions (Figure 3A, lanes: (C+) and Ctrl) in which the apical loop of pre-miR-181a-1 was not shielded and thus was accessible for putative negative regulators that induced the cleavage within the unprotected region of the precursor. As a consequence, in these control reactions, we observed abundant accumulation of pre-miR-181a-1 halves and only low levels of miRs-181a-5p (Figure 3A). The generation of pre-miRNA halves could be a part of mechanisms regulating the differential expression of miRNA families in *X. laevis*. The existence of such transcriptional and/or post-transcriptional regulation has been already suggested by our RNA-seq data showing that in *X. laevis* axons the level of mature miR-181a-1-3p is much lower than miR-181a-2-3p [49].

**Figure 3.**
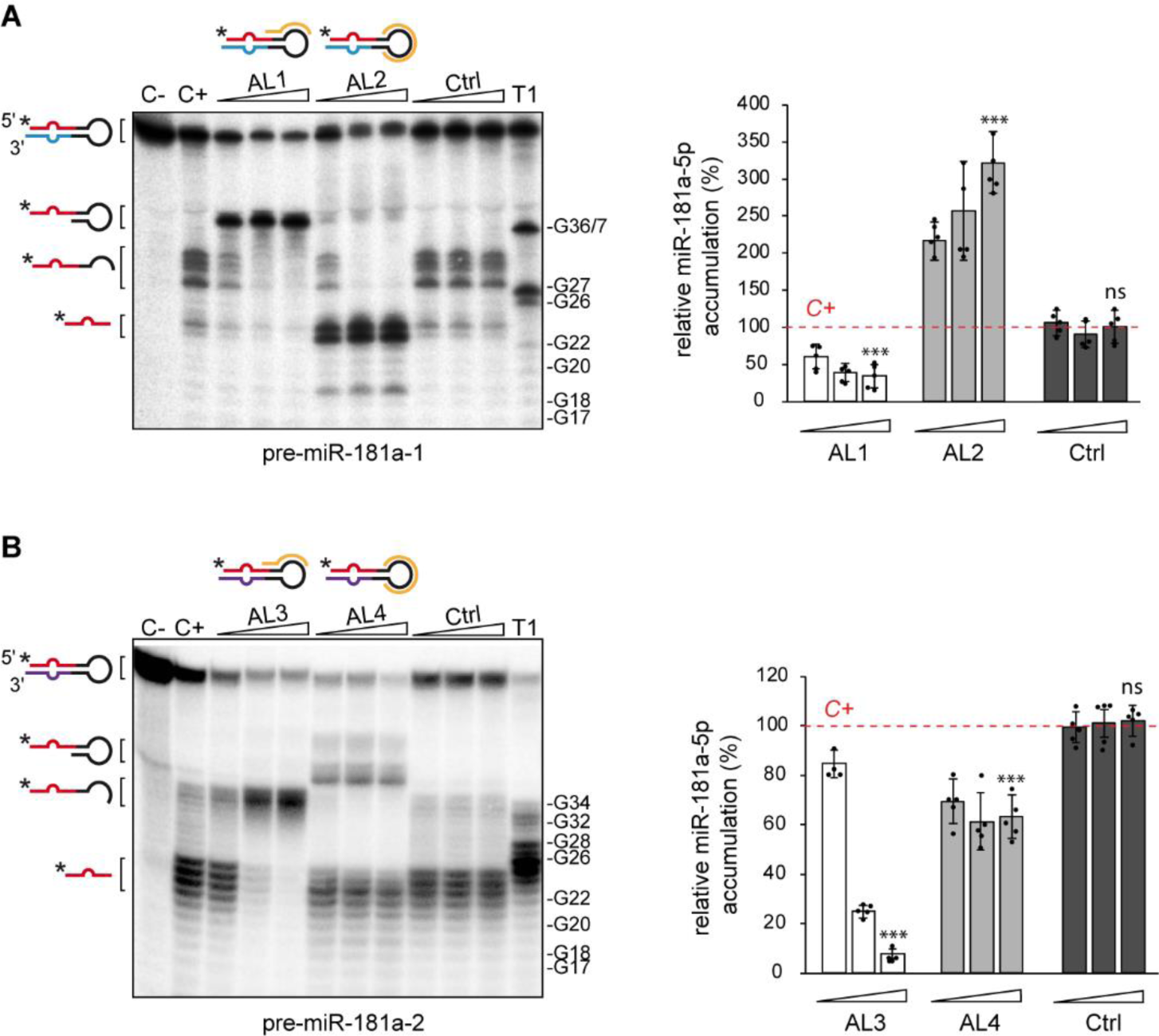
*In vitro* validation of 2′-OMe/LNA ASOs targeting miR-181a-5p precursors. **(A, B)** Results of the inhibition assay assessing the effect of tested oligomers (AL1-4, Ctrl) on the cleavage of 5′-^32^P-labeled pre-miR-181a-1 (A) or 5′-^32^P-labeled pre-miR-181a-2 (B) in *Xenopus* cytosolic extracts. Triangles indicate increasing concentration of a given oligomer (0.1, 1, 10 µM). Graphs show miRNA production efficiency normalized to the level of miRNA generated in C+ for pre-miR-181a-1 (A) and pre-miR-181a-2 (B). Data information: Values are mean ± SD. Statistics: n=5 independent experiments, each data point represents a single assay, unpaired t-test. Abbreviations: ns, not significant; C-, pre-miRNA incubated with no extract nor oligomer added; C+, pre-miRNA incubated with extract but no oligomer added; T1, RNase T1 ladder.

AL3 and AL4, both targeting pre-miR-181a-2, inhibited the formation of miR-181a-5p from this precursor in a dose-response manner, as revealed by the decrease in the intensity or absence of <26-nt bands (Figure 3B, Supplementary Figure S4C, D). However, they did not have the same efficacy in inhibition. Gel densitometry revealed that in reactions with the highest concentration of AL (10 µM), the level of miR-181a-5p was reduced by ∼90% when using AL3 and only by ∼45% when using AL4, in comparison to the positive control reaction (C+). The difference in the inhibitory potential between AL3 and AL4 may be explained by more efficient binding by AL3, as compared to AL4, to pre-miR-181a-2 (Figure 1E), and may also reflect the general higher inhibitory potential of the ASOs overlapping the Dicer cleavage site in comparison to the ASOs complementary only to the apical loop [69]. Additionally, the analysis of RNA conformers by native polyacrylamide gel showed that the 18-nt AL4 can adopt more than one structural form, presumably a homo-duplex or a hairpin (Supplementary Figure S5), which may limit its ability to base-pair with the pre-miRNA and effectively reduce the amount of AL4 available to bind to pre-miR-181a-2 in the reaction. In contrast, 13-nt AL3, AL1, and AL2 adopted only one, monomolecular form (Supplementary Figure S5). These results further indicate that 13-nt ASOs might be more competent inhibitors than longer oligonucleotides, which have a higher possibility of adopting the secondary structures thereby precluding efficient hybridization to the target.

The control oligomer (Ctrl) did not affect miR-181a-5p production efficiency nor pre-miR-181a-1/a-2 cleavage patterns when compared to the positive control reaction (Figure 3A, B). These data and the above results of the native gel electrophoresis (Figure 1E) further affirm that Ctrl represented a suitable control oligomer for further studies.

Taking into consideration the sequence similarity between pre-miR-181a-1 and pre-miR-181a-2, we also performed cross-reactions using pre-miR-181a-2 as a substrate in reactions with AL1-2, and pre-miR-181a-1 in reactions with AL3-4. Under the tested conditions, neither of the oligonucleotides caused any changes in pre-miRNA cleavage efficiencies or patterns, in comparison to the positive control reactions with no inhibitor added (C+) (Supplementary Figure S6A, B). This further strongly indicates that the designed AL-ASOs can discriminate between pre-miRNAs displaying high sequence identity.

Collectively, these data show that oligomers overlapping Dicer cleavage site and the apical region of the precursor, i.e. AL1 and AL3, are more potent inhibitors of the formation of miR-181a-1-5p *in vitro* than oligomers complementary exclusively to the apical loop of the precursor, i.e. AL2 and AL4. These observations made for the *X. laevis* cytosolic extracts are in line with the results previously reported for recombinant human Dicer, which indicate that the inhibitory potential of ASOs can be improved by targeting the Dicer cleavage site [69, 70].

### *In vivo* evaluation of the designed oligomers

#### Assessing inhibitor efficiency in vivo

We next investigated whether 13-nt 2′-OMe/LNA ASOs can be efficiently used in *X. laevis in vivo* at the organismal level. We previously reported successful downregulation of miR-181a-5p level in *X. laevis* using a mix of 24-25-nt MOs targeting either the 5p (MOs-5p) or the 3p (MOs-3p) portion (including the corresponding arm and the apical loop) of both pre-miR-181a-1 and pre-miR-181a-2 [5] (Supplementary Figure S1C-D). Separately, either MOs-5p or MOs-3p blocked the processing of both precursors, simultaneously reducing the levels of miR-181a-5p, miR-181a-1-3p, and miR-181a-2-3p [5]. Here, we investigated whether the robust knockdown of miR-181-5p could be achieved *in vivo* using our shorter 2′-OMe/LNA modified ASOs with precursor specificity. Considering our *in vitro* data (Figure 3), for *in vivo* studies we chose AL1 and AL3 targeting the Dicer cleavage site within the 5′ arms of pre-miR-181a-1 and pre-miR-181a-2, respectively. We started with microinjection of 1 ng of the mixture of AL1 and AL3, or Ctrl oligomer, together with a GFP-expressing plasmid, into dorsal blastomeres at the 8-cell stage (Figure 4A). The targeted cells are fated to form the entire CNS, including the retina [71]. We evaluated the possible toxic effects of the oligomers on *X. laevis* by assessing the survival rate of the embryos following the microinjection. The survival rate of embryos microinjected with either PBS or ALs was similar to WT embryos, highlighting the absence of cytotoxicity of ALs inhibitors in the whole animal (Figure 4B). At stage 20, embryos successfully microinjected and thus expressing GFP in the CNS were selected and grown till stage 40, matching the previous experimental conditions [5]. At stage 40, the eyes were dissected for RNA extraction and analysis by RT-qPCR of the levels of the miRNAs miR-181a-5p, miR-181a-1-3p, miR-181a-2-3p and miR-182 (Figure 4A). miR-182 is also highly expressed in the retina, including in RGC axons [49] and thus, we used it as an internal control for the RT-qPCR reaction. We first measured the expression levels of all four miRNAs upon Ctrl oligomer microinjection: no changes were found comparing Ctrl and WT samples (Supplementary Figure S7), again confirming the suitability of the Ctrl oligomer as a negative control in further experiments. Then, we evaluated the influence of AL1 and AL3 on miR-181a-5p production. Compared to Ctrl, the mix of AL1 and AL3 led to a significant decrease in miR-181a-5p levels (∼72%), while neither the expression level of both miR-181-3p forms nor miR-182 was significantly impacted (Figure 4C). This result differs from what we previously observed with MOs targeting miR-181a-1/-2, where both miR-181-5p and miR-181-3p forms were robustly knocked down [5]. However, similarly to what was seen with the MOs, the mix of AL-ASOs here mediated a long-lasting downregulation of the miR-181-5p level observed even after 8 days post-microinjection (Figure 4C). Together, these data indicate that targeting Dicer 5′ cleavage site with short 2′-OMe/LNA ASOs enables efficient and specific knockdown of miRNA-5p and not miRNA-3p, whereas longer ASOs of different chemistry, such as MOs, can be used to knockdown both miRNAs with a single oligomer.

**Figure 4.**
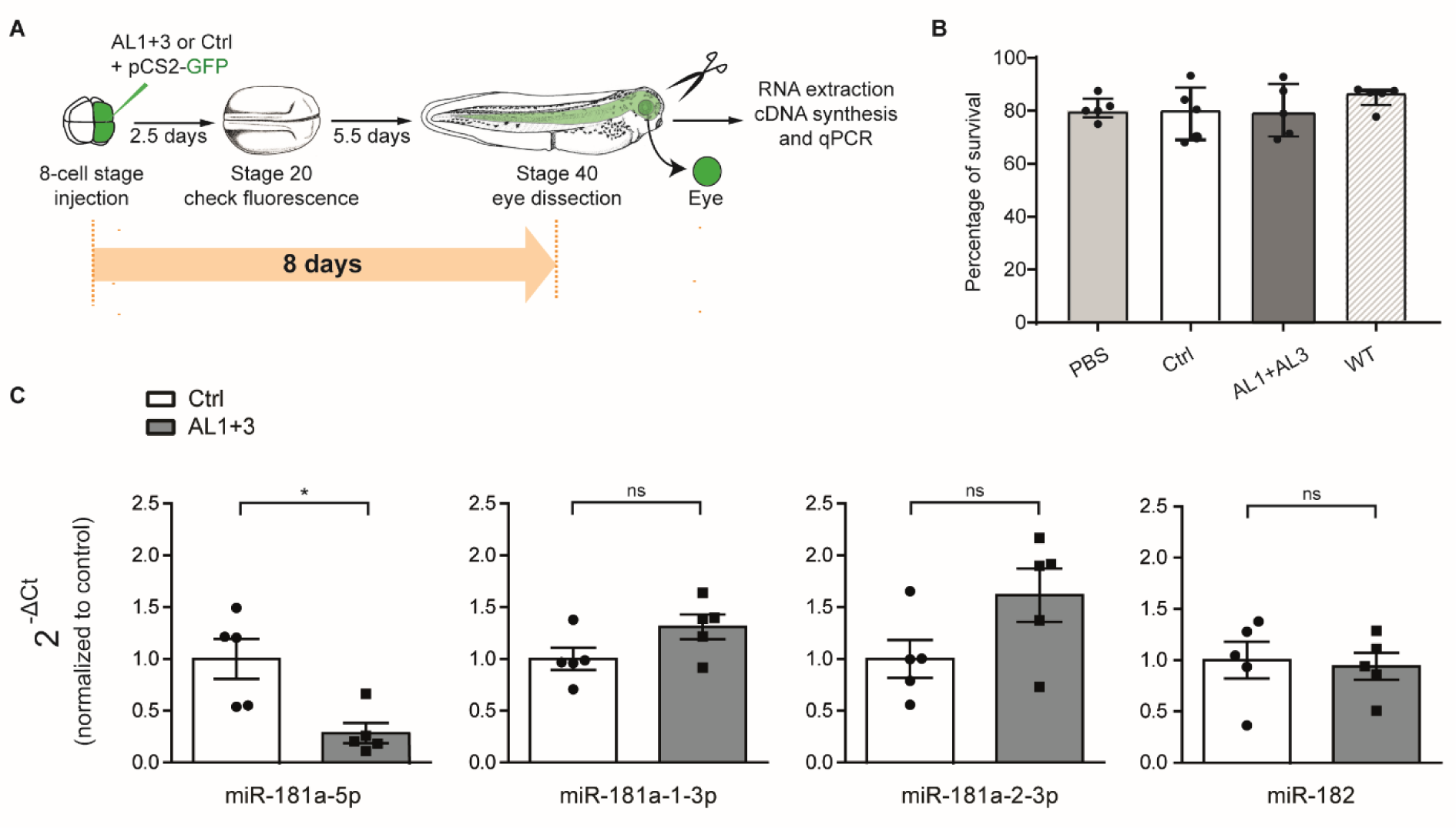
*In vivo* validation of 2′-OMe/LNA ASOs targeting miR-181a-5p precursors using microinjection. **(A)** Schematic of the experimental paradigm. Quantity used: 0.5 ng pCS2-eGFP and 1 ng AL1 and AL3 (anti-181 mix) or 1 ng of the control oligomer (Ctrl). **(B)** Survival embryos rate after microinjection of oligomers (AL1+3, Ctrl) or PBS in comparison to non-injected embryos (WT). **(C)** miRNAs expression levels quantified using the 2^-ΔCt^ method and U6 as normalizer. Data are normalized to PBS control. Data information: Values are mean ± SEM. Statistics: n=5 independent experiments, each data point represents a single RT–qPCR, unpaired t-test. Abbreviations: ns, not significant.

One of the applications of miRNA inhibitors is to study miRNA endogenous function through the miRNA LOF. For such an application, it is important to be able (i) to selectively deliver the inhibitors to the region of interest and (ii) to discriminate *a bona fide* on-target phenotype from possible off-target and toxic effects induced by tools used for the LOF. We examined both aspects using our AL-ASOs. Microinjection is a powerful tool to efficiently deliver the inhibitors at the very early stage of development, into two of the four dorsal blastomere cells that are the source of all descendant cells forming the CNS. This method, however, does not allow for selective targeting of any subset of cells within the CNS (e.g. the retina). To specifically target the retina, we electroporated the AL1 and AL3 mix or the Ctrl oligomer into the retinal primordium [72] using GFP plasmid as a delivery control (Figure 5A). Following the electroporation, embryos did not develop any global malformations and all of them survived after oligomers delivery at stage 25 (Supplementary Figure S8A). Next, we verified whether the oligomers could impair eye development in *X. laevis*. Despite the crucial role of miR-181-5p in RGC axon guidance [5], the knockdown of this miRNA in *X. laevis* does not affect the global eye structure as we previously observed [5] and as also shown for Medaka fish [73]. After the delivery of the mix of AL1 and AL3 into *X. laevis* eye primordia (stage 25), we measured the eye size at stage 40 (Figure 5A), i.e. the stage at which we also evaluated the AL-ASOs efficiency *in vitro* (Figure 3). AL1 and AL3 did not cause any global morphological alterations in the eye (Figure 5B-D). No differences in the eye area (Figure 5C) or diameter (Figure 5D) were observed between electroporated and non-electroporated eyes, at any of the tested oligomer concentrations (up to 500 µM). These data show that electroporation of AL1 and AL3 or Ctrl did not affect the health state of the embryos (Supplementary Figure S8A), nor the development of the electroporated eye (Figure 5A-D). Combined with our cytotoxicity assessment following microinjection (Figure 4B), this collectively suggests that AL inhibitors do not cause embryo- and organo(ocular)-toxicity.

**Figure 5.**
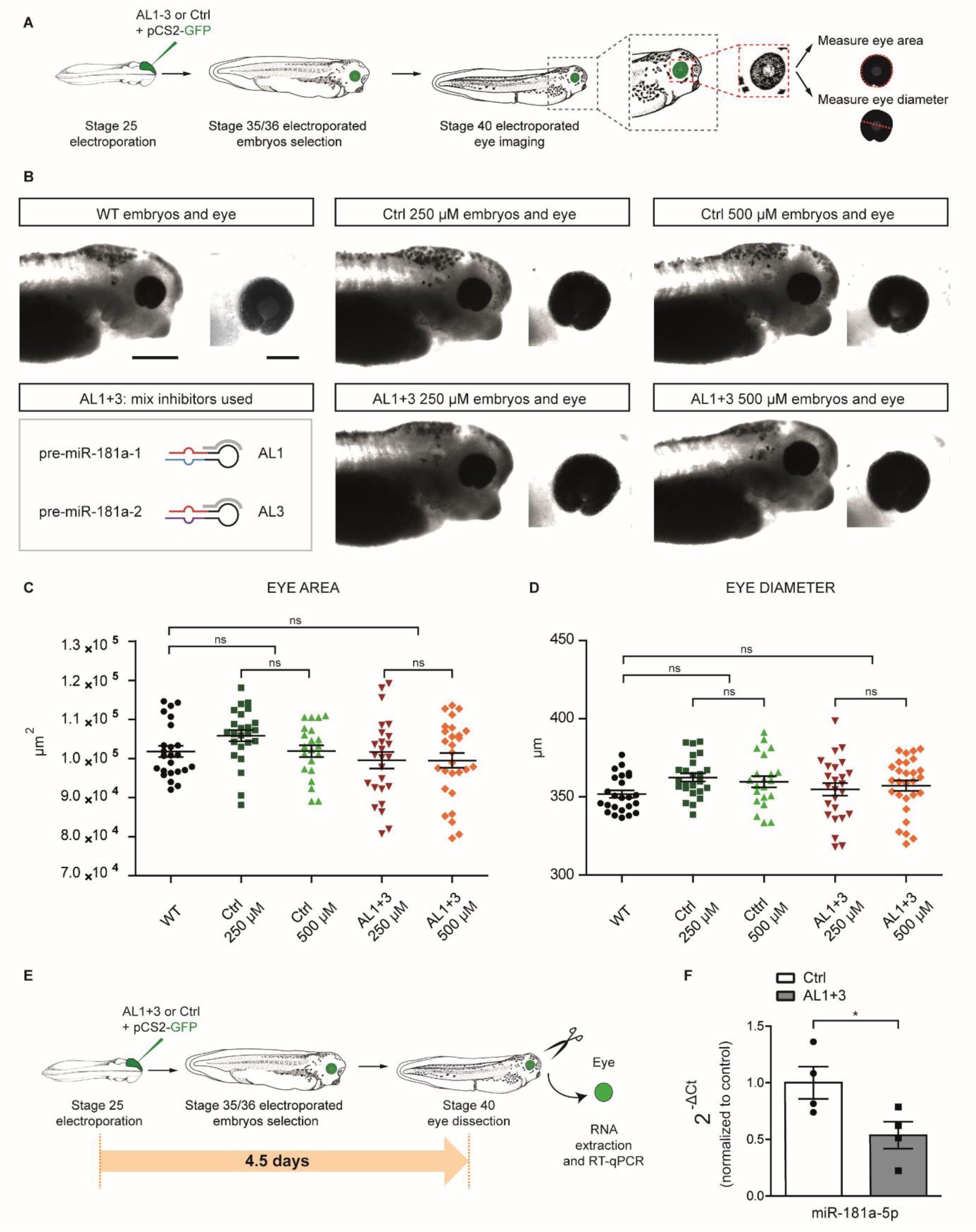
*In vivo* validation of 2′-OMe/LNA ASOs targeting miR-181a-5p precursors using electroporation. **(A,E)** Schematic of the experimental paradigm. Concentrations used: 125 μM anti-181 mix (AL1+AL3) or 125 μM control oligomer (Ctrl), 0.5 μg/μl pCS2-GFP. **(B)** Snapshot of representative electroporated embryos stage 40 in all the experimental conditions (WT, AL1+3 and Ctrl). Bottom left, schematic of the ALs inhibitors used for *in vivo* experiments. **(C, D)** Eye area (C) and diameter (D) of stage 40 wild type, AL1+3 or Ctrl electroporated embryos. **(F)** miR-181a-5p expression levels quantified using the 2^-ΔCt^ method and U6 as normalizer. Data are normalized to PBS control. Each data point represents a single RT–qPCR. Data information: Values are mean ± SEM (C, D, F). Statistics: n=3 independent experiments, 1-way ANOVA, Tukey’s Multiple Comparison Test (C, D); n=4 independent experiments, unpaired t-test (F). Scale bars: 500 μm (B left, embryo), 200 μm (B right, eye). Abbreviations: ns, not significant.

Next, we investigated whether the electroporated inhibitors can efficiently knock down miR-181a-5p expression *in vivo* within the eye. A mix of AL1 and AL3 or Ctrl were electroporated at stage 25, alongside GFP plasmid to label electroporated cells. At stages 35/36 only embryos expressing GFP were selected and miRNA levels were quantified in stage 40 eyes by RT-qPCR (Figure 5E). miR-181a-5p levels decreased (∼46%) upon AL1 and AL3 delivery compared to Ctrl (Figure 5F), while the 3p forms, as well as miR-182, did not change significantly (Supplementary Figure S8B). It should be noted that not all the cells within the eye were successfully electroporated [72]. Therefore, 46% decrease of miR-181-5p level, being the average from a mixed population of targeted and WT cells, represents a good knockdown efficiency. The miRNAs expression levels in Ctrl-treated eyes and WT eyes were comparable (Supplementary Figure S8C).

Altogether these data show that the 13-nt 2′-OMe/LNA ASOs are compatible for use *in vivo* at the organismal level, as they did not cause any sequence- or chemistry-driven off-target effects that could cause developmental malfunctions in *X. laevis* embryos. Additionally, they can be successfully delivered to the cells by at least two popular transfection methods, i.e. microinjection, and electroporation.

#### Intra-axonal AL-ASOs delivery

Recently, we showed that newly generated miR-181-5p molecules in *X. laevis* RGC axons are involved in axon guidance [5]. In particular, we observed that Sema3A, a repellent cue expressed at the boundary of the RGC target region in the brain, induces pre-miR-181a-1/-2 maturation and that the Sema3A response is impaired by blocking Dicer cleavage of these pre-miRNAs specifically in axons [5]. Therefore, we wanted to assess whether this altered axonal Sema3A-response could be reproduced with AL1 and AL3 oligomers. To achieve specific axonal inhibition of the formation of newly-generated miRNAs, and avoid miRNAs contribution from the soma, axons were manually severed from the explant before transfection (Figure 6A, B). After axon isolation, pre-miR-181a-1/-2 were targeted by local transfection of AL1 and AL3 mix using magnetic nanoparticles (Figure 6A, and see Methods for details), as before for MOs [5]. During development, the growth cone, the dynamic tip of a growing axon, promptly responds to the environment, moving towards attractive cues and away from repellent ones. *Ex vivo*, growth cones collapse upon repellent cue exposure, assuming a “tulip shape”, and they can be counted from the total number of growth cones on the culture plate to provide a quantitative measure of growth cone responsiveness to a cue [74]. Here, we used the percentage of collapsed axons as the readout of Sema3A responsiveness. Isolated axons transfected with Ctrl oligomer showed ∼62% collapsed growth cones upon Sema3A bath application, as expected in control conditions [5, 74], while only ∼42% of the axons transfected with the mixture of AL1 and AL3 properly responded to Sema3A (Figure 6C). Moreover, no significant difference was observed between AL1- and AL3-transfected axons upon exposure to Sema3A or PBS (Figure 6C). This indicates that AL1- and AL3-abolished growth cone response to Sema3A. It further suggests that pre-miR-181a-1 and pre-miR-181a-2 processing is essential for Sema-3-mediated response. Such results are similar to those previously obtained using MO oligomers, supporting a critical role of miR-181a-5p in *X. laevis* axon guidance [5]. However, the applied MO oligomers in contrast to AL-ASOs tested here, also reduced the level of miRNA-3p forms: miR-181a-1-3p and miR-181a-2-3p. Since AL-ASOs downregulated selectively only the miR-181-5p levels (Figure 3-5), the collected data for the first time demonstrate the sole importance of miR-181-5p (i.e. not in connection with its 3p counterparts) for the local regulation of the neuronal circuits.

**Figure 6.**
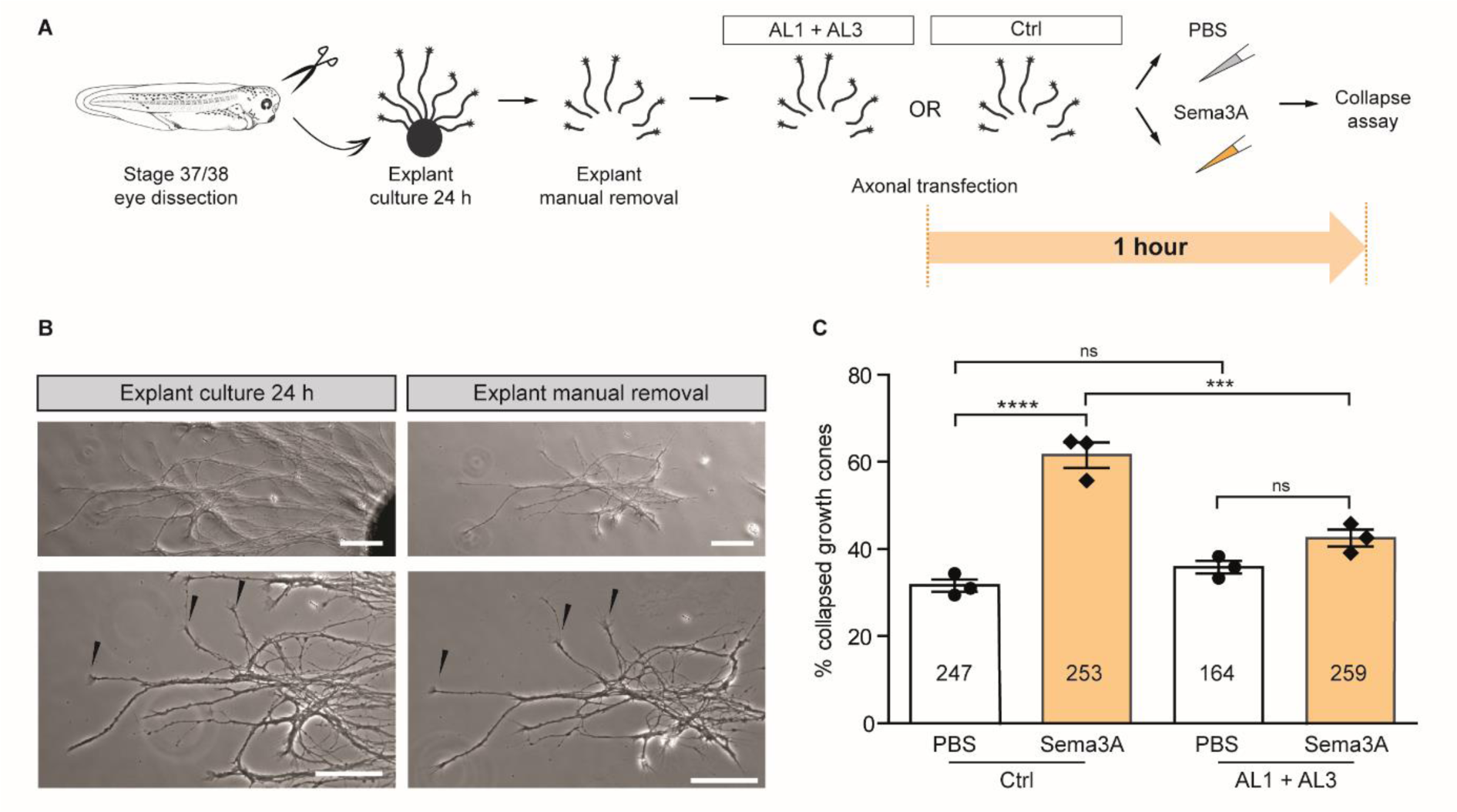
*In vivo* validation 2′-OMe/LNA ASOs targeting miR-181a-5p precursors within axons. **(A)** Schematics of the experimental paradigm. Concentrations used: 2 μM AL1+AL3 or 2 μM Ctrl, 200 ng/mL (Sema3A). **(B)** Representative RGC culture before and after explant manual removal. Severing axons do not cause collapse per se (arrowheads indicate not-collapsed growth cones). **(C)** Frequency (in percentage) of collapsed growth cones after 10 min Sema3A bath application. The number of counted growth cones is reported in each column. Data information: Values are mean ± SEM. Statistics: Each data point corresponds to one independent experiment, n=3 independent experiments, 2-way ANOVA, Sidak’s multiple comparison test. Scale bars: 50 μm. Abbreviations: ns, not significant.

Collectively, our data demonstrate that the tested AL-ASOs are suitable tools for pre-miRNAs inhibition in cellular subcompartments upon transfection, and highlight the efficiency of AL1 and AL3 in inhibition of miR-181-5p maturation *ex vivo* including in axons. Since axons represent a very fragile and delicate subcellular culture system, it also reinforces the notion that tested ALs do not generate cytotoxicity. Altogether, we show that the presented strategy based on 13-nt 2′-OMe /LNA ASOs (i) is compatible with vertebrate models (i.e. no cytotoxic effects); (ii) can be applied using two common delivery methods: microinjection and electroporation, (iii) leads to the efficient reduction of the level of a specific miRNA. Moreover, by recapitulating the impaired Sema3A response upon blocking pre-miR-181a-1/-2 maturation in RGC axons, as previously reported using MO [5], we confirm the suitability of these ASOs for functional miRNAs studies *ex vivo* and *in vivo*.

### Comparison of antisense morpholinos and 2ꞌ-O-methyl/LNA ASOs efficacy and selectivity

Finally, we wanted to assess the knockdown efficacy and selectivity of antisense MO, in comparison to what we have demonstrated for AL-ASO. MOs have become a standard method used to alter gene expression in development in model organisms [75]. In our previous *in vivo* studies [5], 24-25-nt MOs were successfully used to simultaneously inhibit the processing of pre-miR-181a-1 and -181a-2 into mature miRNAs but their selectivity was not examined. Here, we first investigated *in vitro* MOs MO-a1-5p, MO-a1-3p, MO-a2-5p, MO-a2-3p separately and in relation to the individual precursors, similarly to our analysis of AL-ASOs (Figure 3 and Supplementary Figure S4). 25-nt MO-a1-5p and 24-nt MO-a1-3p are complementary to the 5p and 3p arm of pre-miR-181a-1, respectively, and 25-nt MO-a2-5p and 24-nt MO-a2-3p are complementary to the corresponding arms of pre-miR-181a-2 (Figure 7A). Similar to the *in vitro* inhibition assays conducted with AL-ASOs (Figure 3), four MOs were tested individually by using 5′-^32^P-pre-miR-181a-1 (Figure 7B) or 5′-^32^P-pre-miR-181a-2 (Figure 7C), and cytosolic extracts from stage 44 *X. laevis* embryos. The efficiency of 5′-^32^P-miR-181a-5p production was calculated based on the results of PAGE densitometric analysis. MO-a1-5p very efficiently inhibited the formation of miR-181a-5p from pre-miR-181a-1 (Figure 7B, D), but also from pre-miR-181a-2 (Figure 7C, E), since the intensity of miRNA-corresponding bands decreased in a dose-dependent manner compared to the control reaction (C+). Similarly, MO-a2-5p inhibited the production of miR-181a-5p not only from pre-miR-181a-2 (Figure 7C, E) but also from pre-miR-181a-1 (Figure 7B, D). MO-a1-3p affected the cleavage of pre-miR-181a-1 but not pre-miR-181a-2, and MO-a2-3p affected the cleavage of pre-miR-181a-2 but not pre-miR-181a-1 (Figure 7B-E). Consequently, our results indicate that while MOs were very efficient in blocking Dicer-mediated processing of pre-miRNA, neither MO-a1-5p nor MO-a2-5p were precursor-selective. In contrast, both MO-a1-3p and MO-a2-3p discriminated between pre-miR-181a-1 and pre-miR-181a-2. Nevertheless, in both cases, the MO-3p failed to be arm-selective and inhibited not only the release of the miRNA-3p but also the miRNA-5p, since a dose-dependent reduction of the intensity of bands corresponding to the 5′-^32^P-miR-181a-5p isoforms was observed for them (Figure 7B, C). Collectively, these data indicate that MOs targeting Dicer cleavage-site can induce robust knockdown of both mature 3p and 5p miRNAs by interaction with pre-miRNAs precluding Dicer cleavage, but they do not allow to discriminate between highly similar precursors (MO-a1-5p, MO-a2-5p) or to selectively inhibit miRNA only from one pre-miRNA arm (MO-a1-3p, MO-a2-3p).

**Figure 7.**
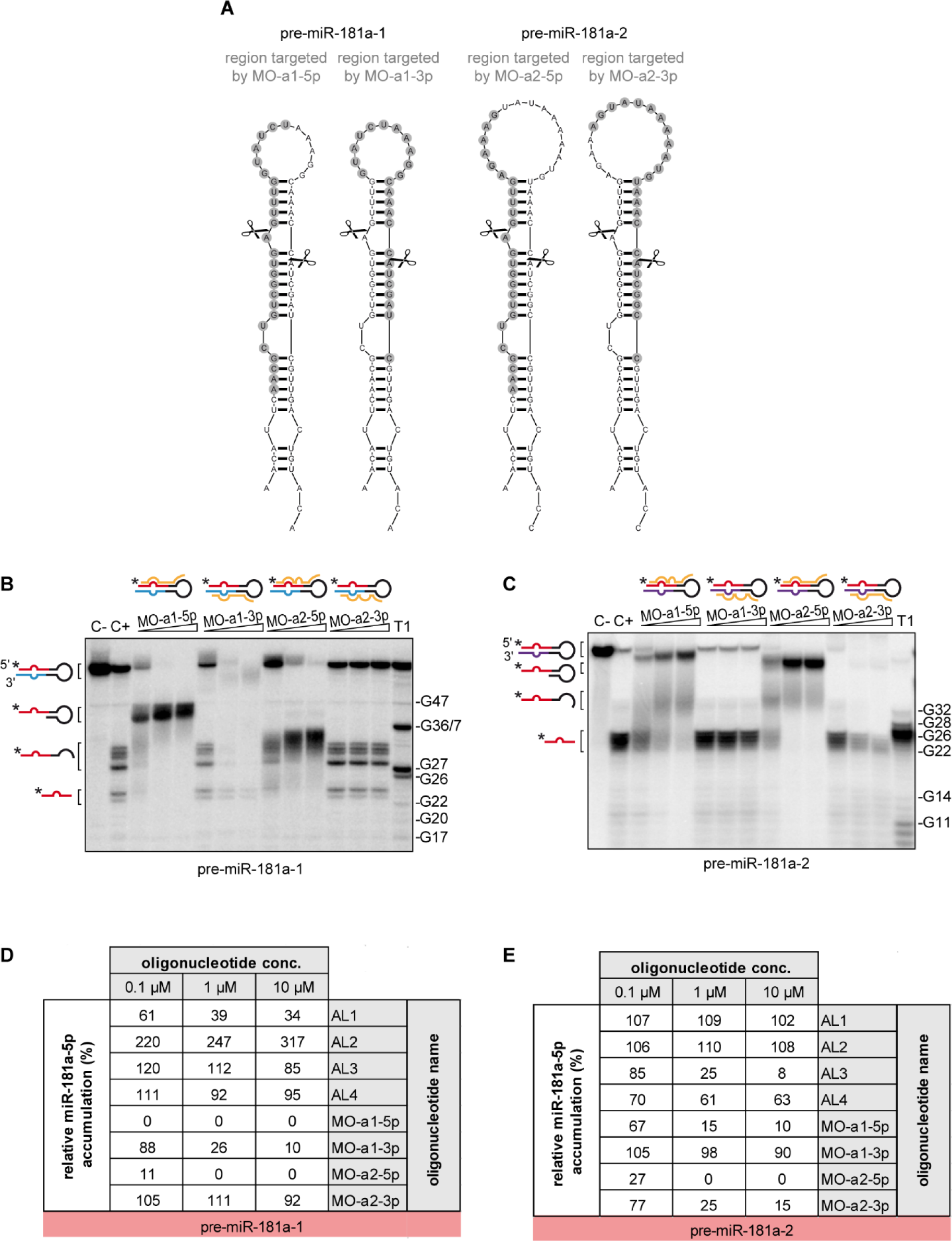
*In vitro* validation of MO ASOs inhibition potential and target-specificity. **(A)** Predicted secondary structures of pre-miR-181a-1 (*left*) and pre-miR-181a-2 (*right*). Scissors mark Dicer cleavage sites. Grey circles indicate the regions targeted by MOs. **(B, C)** Results of the inhibition assay assessing the effect of MOs on the cleavage of 5′-^32^P-labeled pre-miR-181a-1 (B) or 5′-^32^P-labeled pre-miR-181a-2 (C) in *Xenopus* cytosolic extracts. Triangles indicate increasing concentration of a given MO (0.1, 1, 10 µM). Schematic representation of the pre-miRNA hairpin and MO (yellow line) is given above each triangle. **(D, E)** Comparison of the inhibitory potential of AL1-4 and tested MOs calculated as miR-181a-1-5p production efficiency normalized to the level of miR-181a-1-5p generated in C+ for pre-miR-181a-1 (D) and pre-miR-181a-2 (E). Data derived from Fig. 3 and Supplementary Fig. S7 for AL1-4, and (B, C) for MOs. Abbreviations: MO, morpholino; C-, pre-miRNA incubated with no extract nor MO added; C+, pre-miRNA incubated with extract but no MO added; T1, RNase T1 ladder.

It is important to note that the nucleotide sequences of the target regions for 25-nt MO-a1-5p and MO-a2-5p share 80% identity (20/25 nt) (Supplementary Figure S1C), whereas those for 24-nt MO-a1-3p and MO-a2-3p are identical in 75% (18/24 nt) (Supplementary Figure S1D). Since 25-nt MOs-5p proved to be very efficient inhibitors of miR-181a-5p formation [5] (Figure 7B-E), but under the applied reaction conditions failed to discriminate between its precursors, we conclude that the 5-nt difference within the pre-miRNA target region is not sufficient to ensure selectivity for such long ASOs. Beyond this threshold, 25-nt MOs may start to have discriminatory potential as seen for MO-a1-3p (6-nt difference) and MO-a2-3p (7-nt difference). In stark contrast, in the case of 13-nt 2′-OMe/LNA ASOs, despite a smaller, 2-nt difference within the pre-miRNA target region, corresponding to a high sequence similarity, i.e. ∼85% identity (11/13 nt), the high target selectivity is observed (Supplementary Figure S6). Thus, 13-nt 2′-OMe/LNA appears much better suited ASO than 25-nt MO for target-selective knockdown of highly-similar sequences.

Despite years of research and wide interest in the biological roles of miRNAs, only a fraction of known miRNAs has been functionally characterized. This imbalance has many contributing factors. Among these, one critical issue is the outpacing of more laborious functional studies, often hampered by technical limitations, by the availability of next-generation sequencing methods that promote the rapid discovery of new miRNAs. Another key issue is the complication that despite the initial consensus that one pre-miRNA gives rise to only one active regulatory molecule (a guide miRNA), it is now evident that a single precursor can, in fact, encode more than one biologically active entities, including a passenger strand miRNA* [76, 77], loop-miR (miRNA originating from the pre-miRNA loop) [78] or isomiRs (miRNA isoforms) [79]. Accumulating evidence further suggests that the abundance of these additional products of pre-miRNA processing has been underestimated. Our understanding of the physiological relevance of these myriad species and the intricacy of miRNA function has been hampered by current methods that lack the resolution needed for highly selective targeting of only one of these products for functional studies. Moreover, the fact that miRNAs tend to evolve into large and complex families of identical or highly similar members makes it extremely difficult to design antisense tools that would target each of them separately [42] and truly verify the hypothesis that miRNAs within a family have shared or interchangeable functions.

Our novel strategy based on 13-nt long 2′-OMe/LNA ASOs overcomes the limitations of current methods used for miRNA knockdown and allows us to dissect the biological roles of identical or very similar miRNAs originating from different precursors, in cultured cells and *in vivo*. This strategy offers (i) high hybridization affinity due to combining 2′-OMe/LNA modifications, (ii) high target discrimination, which translates to selective targeting of individual, highly similar pre-miRNAs and (iii) long-lasting effects *in vivo* presumably due to AL-ASOs resistance to cellular nucleases. We envision that this new tool will be of critical importance to gain a deeper understanding of the roles of individual miRNAs given the broad significance of miRNAs to physiology and disease.

## ACKNOWLEDGMENTS

We would like to thank Life Science Editors for editing services. This work was supported by the National Science Centre, Poland [2016/22/E/NZ1/00422 to A.K-K.], Marie Curie Career Integration (618969 GUIDANCEmiR), the G. Armenise-Harvard Foundation, MIUR SIR (RBSI144NZ4) and MIUR PRIN 2017 (2017A9MK4R) to M.-L.B.

## METHODS

### OLIGONUCLEOTIDES

Sequences of oligonucleotides used in this study are listed in Supplementary Table S3. 2′-O-methyl/LNA were synthesised in-house and PAGE-purified. Other oligonucleotides were purchased as follows: DNA probes for Northern blot analysis - IBB PAS, pre-miRNA - FUTURESynthesis sp. z o.o., morpholino - GeneTools, all oligonucleotides used for miRNA RT-qPCR quantification - Thermo Fisher Scientific.

### 2′-OMe/LNA OLIGONUCLEOTIDES DESIGN

*X. laevis* mRNA reference sequences were downloaded from RefSeq (1 MM) and UCSC Genome Browser (2 MM) databases. A reference dataset for small RNAs was built using data derived from NGS experiments (GSE86883, GSE33444, PRJNA292052). Oligomers targeting pre-miRNA (pre-miR-181a-1 and pre-miR-181a-2) apical loop region were designed by searching for complementary 13-nt long RNA sequences with the lowest Gibbs free energies of target-oligomer hybridization as calculated using IntaRNA [63] and the lowest potential for the off-target interactions. The latter was tested by the alignment of AL-ASOs reverse-complements to mRNA and small RNA reference sequences using Bowtie [80]. To design a control oligomer, a set of all possible 13-nt long RNA sequences - Potential Target Sites (PTS) - was aligned to either mRNA or small RNA reference dataset using Bowtie. Three rounds of alignment were performed with raising -v parameter (possible number of mismatches, from 0 to 2), each time, unaligned PTS from the previous step were used as input for the next round. The last alignment (-v=2) resulted in no unaligned PTS. One of the input PTS from the last alignment round was selected and converted into reverse-complement to obtain a control oligomer sequence (Ctrl). The self- and cross-hybridization free Gibbs energies of AL1-4 and Ctrl were calculated using EvOligo [64].

### 32P LABELING OF OLIGONUCLEOTIDES

RNA (10 µM) was labeled with 1 µL of [γ^32^P] ATP (3000 Ci/mmol, Hartman Analytic GmbH) and 10 U T4 polynucleotide kinase (Thermo Fisher Scientific) for 10 min at 37°C. The radiolabeled RNA were PAGE-purified in 8% denaturing polyacrylamide gels and resuspended in water to a final concentration of approximately 10,000 cpm/µL. DNA probes (10 µM) for Northern blot were labeled with 5 µL [γ^32^P] ATP (5000 Ci/mmol; Hartmann Analytics) and 10 U T4 Polynucleotide Kinase (Thermo Fisher Scientific) for 1 h at 4°C. Probes were purified with NucAway spin columns (Ambion), an entire 50 µL elution was used to probe a single membrane.

### RNA NATIVE GEL ELECTROPHORESIS

RNA-RNA interaction was tested as described earlier [69]. Briefly, 5′-^32^P-labeled pre-miRNA (1 pmol, 10 000 cpm per sample) was mixed with 100-molar excess of tested oligomer (AL1-4 or Ctrl) in binding buffer (20 mM Tris-HCl, pH 7.5, 100 mM NaCl), and incubated for 30 min at 22°C. The samples were analyzed by non-denaturing PAGE. Gel imaging was performed using FLA-5100 Fluorescent Image Analyzer (Fujifilm).

### *XENOPUS LAEVIS* EMBRYOS MAINTENANCE

*X. laevis* embryos were obtained by *in vitro* fertilization and kept in autoclaved 0.1x Modified Barth’s saline diluted in double distillate water ddH_2_O pH 7.5 (MBS 10x: 88 mM NaCl, 1mM KCl, 0.82 mM MgSO_4_, 2.4 mM NaHCO_3_, 10 mM HEPES, 0.33 mM Ca(NO_3_)_2_, 0.41 mM CaCl_2_) at 14-22°C. They were staged according to Nieuwkoop and Faber [81]. All animal experiments were approved by the University of Trento Ethical Review Committee and by the Italian “Ministero della Salute” both according to the D.Lgs nr.116/92 and with the authorization n°1159/2016-PR and n°546/2017-PR according to art.31 of D.lgs. 26/2014.

### PREPARATION OF CYTOSOLIC EXTRACTS FROM *XENOPUS LAEVIS* EMBRYOS

For each biological replicate, 25 stage 44 embryos were washed three times in MMR 0.1% and homogenized in 100 μL of ice-cold Evans lysis buffer (1:100 protease inhibitor cocktail (Sigma), 50 mM NaCl, 10 mM MgCl_2_, 20 mM Tris-HCl, pH 7.6). The homogenate was layered on top of a 400 µL ice-cold sucrose cushion (20% sucrose in Evans lysis buffer) in a 1.5 mL microtube and centrifuged at 10000 xg for 30 min at 4°C. The cytosolic fraction (supernatant and a floating phase) was collected and stored at -80°C.

### CLEAVAGE ASSAY

5′-^32^P-labeled pre-miRNA was incubated in *Xenopus* cytosolic extract (10 µg of total protein) with RNase inhibitor (10U of RNaseOUT, Invitrogen) and carrier RNA (2 µg of yeast tRNA, Thermo Fisher Scientific). Samples were incubated for 1 h at 22°C unless stated otherwise in the figure legend. Reactions with human recombinant Dicer (hDicer) were incubated for 30 min at 37°C in a standard cleavage buffer (20 mM Tris-HCl pH 7.5, 50 mM NaCl, 2.5 mM MgCl_2_). The reactions were stopped by the addition of 1 volume of urea loading buffer and heating at 95°C for 5 min, and analysed by denaturing PAGE (15% PAA with 7 M urea) in 1x TBE. Data were collected using Fujifilm FLA-5100 Fluorescent Image Analyzer and analysed using MultiGauge 3.0 (Fujifilm).

### INHIBITION ASSAY

5′-^32^P-labeled pre-miRNA was incubated with the indicated oligonucleotide (0.1 µM, 1 µM, 10 µM) in *Xenopus* cytosolic extract (10 µg of total protein) supplemented with RNase inhibitor (10U of RNaseOUT, Invitrogen), and carrier RNA (2 µg of yeast tRNA, Thermo Fisher Scientific). In addition, two control reactions were carried out: (i) a negative control (*C-*) with no *Xenopus* cytosolic extract and no inhibitor, to test the integrity of the substrate during the incubation time, and (ii) a positive control (*C+*) with *Xenopus* cytosolic extract but no inhibitor added. All samples were incubated for 1 h at 22°C. The reactions were stopped by addition of 1 volume of urea loading buffer and heating for 5 min at 95°C, and analysed by denaturing PAGE (15% PAA with 7 M urea) in 1x TBE. The amounts of radiolabeled pre-miRNA and cleavage product/s in each reaction were quantified by gel densitometry. Data were collected using Fujifilm FLA-5100 Fluorescent Image Analyzer and analysed using MultiGauge 3.0 (Fujifilm). The efficiency of miRNA production in the presence or absence of the oligomer was calculated to determine the oligomers’ capacity to inhibit pre-miRNA digestion by Dicer. The influence of the oligomers on miRNA production was expressed as a percentage, with the miRNA production in control reactions lacking the oligomer (C+) defined as 100%.

### *XENOPUS LAEVIS* EYE ELECTROPORATION

RGC electroporation was performed as previously described [72]. In particular, stage 25 embryos were anesthetized in 0.3 mg/mL MS222 (Sigma) in 1x MBS. The nucleic acid mixture was injected in retinal primordium using a 1.0 mm outer diameter (OD) x 0.5 mm or 0.78 mm internal diameter (ID) glass capillary (Harvard Apparatus). Next, the injected mixture was delivered into the cells at 18 V, by applying 8 electric pulses of 50 ms duration at 1000 ms intervals. After the electroporation, the embryos were rinsed in 0.1x MMR and grown until the developmental stage of interest.

### *XENOPUS LAEVIS* MICROINJECTION

4-cells embryos were dejelled into a 0.2 M Tris (Ambion) and 0.2 M DTT (Thermo Fisher Scientific) water solution for a few minutes and washed in 0.1x MMR. At the 8-cells stage, embryos were placed into an injection dish containing 4% Ficoll (Carl RothGmBH). 1 ng of inhibitors mixture (AL1+AL3) or control oligomer and 0.5 ng of pCS2-eGFP plasmid were microinjected into both animal blastomeres using a 1.0 mm outer diameter (OD) x 0.5 mm internal diameter (ID) glass capillary injection needle (Harvard Apparatus). At stage 19 embryos were selected based on GFP expression in the entire central nervous system as a proxy of successful AL inhibitors delivery, and raised until stage 40 for eye dissection and RNA extraction.

### RETINAL EXPLANTS CULTURING

Glass-bottom dishes (MatTek) were coated with poly-L-lysine (Sigma, 10 μg/mL diluted in double distilled water (ddH_2_O)) for 3 h, washed three times with ddH_2_O, and dried for 10 minutes. Dishes were then coated with laminin (Sigma) 10 μg/mL, diluted in L-15 medium (Gibco), for 1 h at room temperature, followed by two washes with complete culture medium (60% L-15 in ddH_2_O, supplemented with 1% Antibiotic-Antimycotic (Thermo Fisher Scientific).

Before dissection, embryos were washed three times in 0.1% MMR (10x MMR in ddH_2_O and 1% Antibiotic-Antimycotic) and then anesthetized with 0.3 mg/mL MS222 (60% L-15 in ddH_2_O, 1% Antibiotic-Antimycotic and MS222 (Sigma), pH 7.6-7.8). Anesthetized stage 37/38 embryos were secured laterally with custom-made pins on a sylgard dish. Both eyes of wild-type embryos or treated eyes in the case of electroporated embryos were dissected, washed twice in 60% L-15, plated on the pre-coated dishes containing culture medium, and cultured at 20°C for 24 hours in 60% L-15 and 1% Antibiotic-Antimycotic.

### ISOLATION AND TRANSFECTION OF AXONS

Retinal ganglion cells (RGC) axons were isolated from stage 37/38 retinal explants cultured for 24 h as described above. The explants were manually removed at the stereomicroscope using two pins (0.20 mm). All explants were removed from the plate using a p10 pipette without perturbing the severed axons. The complete experimental procedure was concluded within 1.5 h after the first cut was made.

For the axonal transfection, AL1 and AL3 suspended in the culture medium were mixed 1:100 with NeuroMag Transfection Reagent (OZ Biosciences) and incubated for 20 min at room temperature. The transfection mixture was then applied to the plate with freshly isolated axons at 2 μM final concentration. The culture dishes were incubated for 15 min at room temperature on the magnetic base (OZ Biosciences) according to the manufacturer’s instructions. The plates were then removed from the magnetic base, incubated for 40 min at 20°C, and washed five times with 200 µL of the complete culture medium without perturbing the axons in the plate.

### COLLAPSE ASSAY

Axons transfected with the mixture of AL1 and AL3, or a control oligomer (Ctrl) were bathed in dose protein-synthesis dependent Sema3A (R&D System) concentration (200 ng/ml) [5], or PBS (for control) for 10 minutes. Axons were then fixed in 2% paraformaldehyde (PFA, Life Technologies) and 7.5% (wt/vol) sucrose (ACS reagent) for 30 minutes, washed three times with 400 µL 1x PBS. Samples were visually inspected under the light microscope. To avoid subjective bias, the analysis was performed in a blind fashion. Growth cones were considered collapsed when they possessed either no filopodia or one or two filopodia, each shorter than 10 μm [74]. A minimal length of 100 μm from the cut was checked. The experiment was run in three independent replicates, each time 14 explants were cultured per condition. The percentage of collapsed growth cones on the total number of the counted ones was plotted and compared among conditions with a 2-way ANOVA multiple comparison test (Prism).

### RNA EXTRACTION AND qPCR ANALYSIS

Total RNA from dissected stage 40 eyes was extracted using Norgen Total RNA Purification Micro Kit (Norgen). 200 µL of the lysis buffer (Buffer RL, Norgen) was added to the eyes and incubated for 5 minutes. The lysate was transferred to a new tube and 120 µL of absolute ethanol (Sigma) was added and mixed by vortexing for 10 seconds. RNA binding to columns and column washing steps were performed following the manufacturer’s instructions and an on-column RNAse free-DNAse I treatment (Norgen) was run. RNA was eluted by applying 18 µL Elution Solution to the column, collected in low binding RNAse-free tubes, and stored at -80°C.

miRNA was quantified using TaqMan™ MicroRNA Assay (4427975 or 4440886). 10 ng of total RNA isolated from electroporated eyes was retrotranscribed in a gene-specific manner using the TaqMan™ MicroRNA Reverse Transcription Kit (Thermo Fisher Scientific) and the TaqMan qPCR assay (Thermo Fisher Scientific) in a reaction volume of 15 µL following the manufacturer’s instructions. The obtained cDNA was diluted 1:5 in DNAse/RNase-free water (Sigma) and 3 µL of it was used as input in the qPCR reaction. qPCR was carried out using TaqMan™ Universal Master Mix II (Thermo Fisher Scientific) and CFX96 detection system (Bio-Rad). Primers used: miR-181a-5p (ID: 000480); miR-181a-1-3p (ID: 004367_mat); miR-181a-2-3p (ID:005555_mat); miR-182 (ID: 000597) and snU6 (ID: 001973).

For RT-qPCR quantitative analysis, cycle threshold (Ct) was defined with CFX96 Bio-Rad software v3.1, as the mean of three technical replicates characterized by a standard deviation smaller than 0.35. MiRNA expression levels were investigated with the ΔCt method [82] applied as follows: 1/(2^[(Ct_miRX_-Ct_U6_)]). Each experiment was run for at least three independent biological samples (the number of independent experiments is indicated in Figure).

### MEASUREMENT OF *XENOPUS LAEVIS* EYES AREA AND DIAMETER

*X. laevis* embryos were anesthetized with 0.3 mg/mL MS222 (Sigma) in MMR 0.1%, pH 7.6-7.8. Electroporated embryos were selected based on pCS2-eGFP expression and their eyes were imaged with Leica DMi8 epifluorescent microscope coupled with a CMOS monochromatic camera (AndorZyla 4.2 CL10-VSC00962, 4,2 Megapixel). The acquisition mode was set to 12-bit grayscale dark field. No binning was applied to the acquisition. Exposure time and light intensity were chosen to optimize the visualization of the eye in whole embryos. The eye area and diameter were measured in ImageJ using respectively the oval and straight selection tools. One-way ANOVA followed by Tukey’s multiple comparison post-hoc test was run to investigate the difference among conditions.

### NORTHERN BLOT

RNA aliquots (1 µM) obtained in cleavage reactions *using Xenopus* cytosolic extracts were resolved by denaturing PAGE (20% PAA, 7M urea) in 0.5x TBE. RNA was transferred to a Hybond N+ hybridization membrane (Amersham) using semi-dry electroblotting (Bio-Rad), cross-linked with UV irradiation (120 mJ/cm^2^) (UVP), and baked in an oven for 30 min at 80°C. The membranes were probed with specific DNA oligonucleotides targeting the 3′ or 5′ arm of the precursor (Supplementary Table S3). The hybridization was carried out overnight at 37°C in a buffer containing 5x SSC, 1% SDS, and 1x Denhardt’s solution. Radioactive signals were visualized by phosphorimaging (FLA-5100 Fluorescent Image Analyzer; Fujifilm). Next, the membrane was stripped by washing in boiling-hot 0.1% SDS three times for 15 min. The efficiency of the stripping was verified by phosphorimaging, and the membrane was re-probed with another DNA oligonucleotide.

### STATISTICS

All data were analyzed with Prism (GraphPad 6 or 7) and all experiments were performed in at least three independent biological replicates. For all tests, the significance level was α = 0.05. The exact number of replicates, tests used, and statistics are reported in the Figure legend. *p≤0.05, **p≤0.01, ***p≤0.001, ****p≤0.0001, ns: non-significant.

## Notes

### Competing Interest Statement

The authors have declared no competing interest.

## REFERENCES

1. Bartel DP. Metazoan MicroRNAs. Cell. 2018 Mar 22;173(1):20–51.

2. Pasquinelli AE, Ruvkun G. Control of developmental timing by micrornas and their targets. Annu Rev Cell Dev Biol. 2002;18:495–513.

3. Wienholds E, Plasterk RH. MicroRNA function in animal development. FEBS Lett. 2005 Oct 31;579(26):5911–22.

4. Chawla G, Sokol NS. MicroRNAs in Drosophila development. Int Rev Cell Mol Biol. 2011;286:1–65.

5. Corradi E, Dalla Costa I, Gavoci A, Iyer A, Roccuzzo M, Otto TA, et al. Axonal precursor miRNAs hitchhike on endosomes and locally regulate the development of neural circuits. EMBO J. 2020 Mar 16;39(6):e102513.

6. Iyer AN, Bellon A, Baudet ML. microRNAs in axon guidance. Front Cell Neurosci. 2014;8:78.

7. Bi Y, Liu G, Yang R. MicroRNAs: novel regulators during the immune response. J Cell Physiol. 2009 Mar;218(3):467–72.

8. Scaria V, Hariharan M, Maiti S, Pillai B, Brahmachari SK. Host-virus interaction: a new role for microRNAs. Retrovirology. 2006 Oct 11;3:68.

9. Sullivan CS, Ganem D. MicroRNAs and viral infection. Mol Cell. 2005 Oct 7;20(1):3–7.

10. McManus MT. MicroRNAs and cancer. Semin Cancer Biol. 2003 Aug;13(4):253–8.

11. Sun E, Shi Y. MicroRNAs: Small molecules with big roles in neurodevelopment and diseases. Exp Neurol. 2015 Jun;268:46–53.

12. Tan G, Tang X, Tang F. The role of microRNAs in nasopharyngeal carcinoma. Tumour Biol. 2015 Jan;36(1):69–79.

13. Condrat CE, Thompson DC, Barbu MG, Bugnar OL, Boboc A, Cretoiu D, et al. miRNAs as Biomarkers in Disease: Latest Findings Regarding Their Role in Diagnosis and Prognosis. Cells. 2020 Jan 23;9(2).

14. Rupaimoole R, Slack FJ. MicroRNA therapeutics: towards a new era for the management of cancer and other diseases. Nat Rev Drug Discov. 2017 Mar;16(3):203–22.

15. Treiber T, Treiber N, Plessmann U, Harlander S, Daiss JL, Eichner N, et al. A Compendium of RNA-Binding Proteins that Regulate MicroRNA Biogenesis. Mol Cell. 2017 Apr 20;66(2):270–84 e13.

16. Ameres SL, Martinez J, Schroeder R. Molecular basis for target RNA recognition and cleavage by human RISC. Cell. 2007 Jul 13;130(1):101–12.

17. Okamura K, Phillips MD, Tyler DM, Duan H, Chou YT, Lai EC. The regulatory activity of microRNA* species has substantial influence on microRNA and 3’ UTR evolution. Nat Struct Mol Biol. 2008 Apr;15(4):354–63.

18. Yang JS, Phillips MD, Betel D, Mu P, Ventura A, Siepel AC, et al. Widespread regulatory activity of vertebrate microRNA* species. RNA. 2011 Feb;17(2):312–26.

19. Lewis BP, Burge CB, Bartel DP. Conserved seed pairing, often flanked by adenosines, indicates that thousands of human genes are microRNA targets. Cell. 2005 Jan 14;120(1):15–20.

20. Abbott AL, Alvarez-Saavedra E, Miska EA, Lau NC, Bartel DP, Horvitz HR, et al. The let-7 MicroRNA family members mir-48, mir-84, and mir-241 function together to regulate developmental timing in Caenorhabditis elegans. Dev Cell. 2005 Sep;9(3):403–14.

21. Alvarez-Saavedra E, Horvitz HR. Many families of C. elegans microRNAs are not essential for development or viability. Curr Biol. 2010 Feb 23;20(4):367–73.

22. Brancati G, Grosshans H. An interplay of miRNA abundance and target site architecture determines miRNA activity and specificity. Nucleic Acids Res. 2018 Apr 20;46(7):3259–69.

23. Broughton JP, Lovci MT, Huang JL, Yeo GW, Pasquinelli AE. Pairing beyond the Seed Supports MicroRNA Targeting Specificity. Mol Cell. 2016 Oct 20;64(2):320–33.

24. Moore MJ, Scheel TK, Luna JM, Park CY, Fak JJ, Nishiuchi E, et al. miRNA-target chimeras reveal miRNA 3’-end pairing as a major determinant of Argonaute target specificity. Nat Commun. 2015 Nov 25;6:8864.

25. Brennecke J, Stark A, Russell RB, Cohen SM. Principles of microRNA-target recognition. PLoS Biol. 2005 Mar;3(3):e85.

26. Liu G, Min H, Yue S, Chen CZ. Pre-miRNA loop nucleotides control the distinct activities of mir-181a-1 and mir-181c in early T cell development. PLoS One. 2008;3(10):e3592.

27. Yue SB, Trujillo RD, Tang Y, O’Gorman WE, Chen CZ. Loop nucleotides control primary and mature miRNA function in target recognition and repression. RNA Biol. 2011 Nov-Dec;8(6):1115–23.

28. Trujillo RD, Yue SB, Tang Y, O’Gorman WE, Chen CZ. The potential functions of primary microRNAs in target recognition and repression. EMBO J. 2010 Oct 6;29(19):3272–85.

29. Das S, Kohr M, Dunkerly-Eyring B, Lee DI, Bedja D, Kent OA, et al. Divergent Effects of miR-181 Family Members on Myocardial Function Through Protective Cytosolic and Detrimental Mitochondrial microRNA Targets. J Am Heart Assoc. 2017 Feb 27;6(3).

30. Loughlin FE, Gebert LF, Towbin H, Brunschweiger A, Hall J, Allain FH. Structural basis of pre-let-7 miRNA recognition by the zinc knuckles of pluripotency factor Lin28. Nat Struct Mol Biol. 2012 Dec 11;19(1):84–9.

31. Thornton JE, Chang HM, Piskounova E, Gregory RI. Lin28-mediated control of let-7 microRNA expression by alternative TUTases Zcchc11 (TUT4) and Zcchc6 (TUT7). RNA. 2012 Oct;18(10):1875–85.

32. Viswanathan SR, Daley GQ. Lin28: A microRNA regulator with a macro role. Cell. 2010 Feb 19;140(4):445–9.

33. Kawahara Y, Mieda-Sato A. TDP-43 promotes microRNA biogenesis as a component of the Drosha and Dicer complexes. Proc Natl Acad Sci U S A. 2012 Feb 28;109(9):3347–52.

34. Forstemann K, Horwich MD, Wee L, Tomari Y, Zamore PD. Drosophila microRNAs are sorted into functionally distinct argonaute complexes after production by dicer-1. Cell. 2007 Jul 27;130(2):287–97.

35. Steiner FA, Hoogstrate SW, Okihara KL, Thijssen KL, Ketting RF, Plasterk RH, et al. Structural features of small RNA precursors determine Argonaute loading in Caenorhabditis elegans. Nat Struct Mol Biol. 2007 Oct;14(10):927–33.

36. Su Y, Yuan J, Zhang F, Lei Q, Zhang T, Li K, et al. MicroRNA-181a-5p and microRNA-181a-3p cooperatively restrict vascular inflammation and atherosclerosis. Cell Death Dis. 2019 May 7;10(5):365.

37. Presnell SR, Al-Attar A, Cichocki F, Miller JS, Lutz CT. Human natural killer cell microRNA: differential expression of MIR181A1B1 and MIR181A2B2 genes encoding identical mature microRNAs. Genes Immun. 2014 Jan-Feb;16(1):89–98.

38. Ebert MS, Neilson JR, Sharp PA. MicroRNA sponges: competitive inhibitors of small RNAs in mammalian cells. Nat Methods. 2007 Sep;4(9):721–6.

39. Meister G, Landthaler M, Dorsett Y, Tuschl T. Sequence-specific inhibition of microRNA- and siRNA-induced RNA silencing. RNA. 2004 Mar;10(3):544–50.

40. Orom UA, Kauppinen S, Lund AH. LNA-modified oligonucleotides mediate specific inhibition of microRNA function. Gene. 2006 May 10;372:137–41.

41. Krutzfeldt J, Rajewsky N, Braich R, Rajeev KG, Tuschl T, Manoharan M, et al. Silencing of microRNAs in vivo with ‘antagomirs’. Nature. 2005 Dec 1;438(7068):685–9.

42. Ebert MS, Sharp PA. MicroRNA sponges: progress and possibilities. RNA. 2010 Nov;16(11):2043–50.

43. Chen CZ. An unsolved mystery: the target-recognizing RNA species of microRNA genes. Biochimie. 2013 Sep;95(9):1663–76.

44. Liu Y, Bergan R. Improved intracellular delivery of oligonucleotides by square wave electroporation. Antisense Nucleic Acid Drug Dev. 2001 Feb;11(1):7–14.

45. Nenni MJ, Fisher ME, James-Zorn C, Pells TJ, Ponferrada V, Chu S, et al. Xenbase: Facilitating the Use of Xenopus to Model Human Disease. Front Physiol. 2019;10:154.

46. Wheeler GN, Brandli AW. Simple vertebrate models for chemical genetics and drug discovery screens: lessons from zebrafish and Xenopus. Dev Dyn. 2009 Jun;238(6):1287–308.

47. Blum M, Ott T. Xenopus: An Undervalued Model Organism to Study and Model Human Genetic Disease. Cells Tissues Organs. 2019;205(5-6):303–13.

48. Baudet ML, Zivraj KH, Abreu-Goodger C, Muldal A, Armisen J, Blenkiron C, et al. miR-124 acts through CoREST to control onset of Sema3A sensitivity in navigating retinal growth cones. Nat Neurosci. 2012 Dec 4;15(1):29–38.

49. Bellon A, Iyer A, Bridi S, Lee FCY, Ovando-Vazquez C, Corradi E, et al. miR-182 Regulates Slit2-Mediated Axon Guidance by Modulating the Local Translation of a Specific mRNA. Cell Rep. 2017 Jan 31;18(5):1171–86.

50. Ahmed A, Ward NJ, Moxon S, Lopez-Gomollon S, Viaut C, Tomlinson ML, et al. A Database of microRNA Expression Patterns in Xenopus laevis. PLoS One. 2015;10(10):e0138313.

51. Lund E, Sheets MD, Imboden SB, Dahlberg JE. Limiting Ago protein restricts RNAi and microRNA biogenesis during early development in Xenopus laevis. Genes Dev. 2011 Jun 1;25(11):1121–31.

52. Indrieri A, Carrella S, Carotenuto P, Banfi S, Franco B. The Pervasive Role of the miR-181 Family in Development, Neurodegeneration, and Cancer. Int J Mol Sci. 2020 Mar 18;21(6).

53. Lennox KA, Behlke MA. Chemical modification and design of anti-miRNA oligonucleotides. Gene Ther. 2011 Dec;18(12):1111–20.

54. Woolf TM, Melton DA, Jennings CG. Specificity of antisense oligonucleotides in vivo. Proc Natl Acad Sci U S A. 1992 Aug 15;89(16):7305–9.

55. Obad S, dos Santos CO, Petri A, Heidenblad M, Broom O, Ruse C, et al. Silencing of microRNA families by seed-targeting tiny LNAs. Nat Genet. 2011 Mar 20;43(4):371–8.

56. You Y, Moreira BG, Behlke MA, Owczarzy R. Design of LNA probes that improve mismatch discrimination. Nucleic Acids Res. 2006 May 2;34(8):e60.

57. Owczarzy R, You Y, Groth CL, Tataurov AV. Stability and mismatch discrimination of locked nucleic acid-DNA duplexes. Biochemistry. 2011 Nov 1;50(43):9352–67.

58. Majlessi M, Nelson NC, Becker MM. Advantages of 2’-O-methyl oligoribonucleotide probes for detecting RNA targets. Nucleic Acids Res. 1998 May 1;26(9):2224–9.

59. Cummins LL, Owens SR, Risen LM, Lesnik EA, Freier SM, McGee D, et al. Characterization of fully 2’-modified oligoribonucleotide hetero- and homoduplex hybridization and nuclease sensitivity. Nucleic Acids Res. 1995 Jun 11;23(11):2019–24.

60. Frieden M, Hansen HF, Koch T. Nuclease stability of LNA oligonucleotides and LNA-DNA chimeras. Nucleosides Nucleotides Nucleic Acids. 2003 May-Aug;22(5-8):1041–3.

61. Fratczak A, Kierzek R, Kierzek E. LNA-modified primers drastically improve hybridization to target RNA and reverse transcription. Biochemistry. 2009 Jan 27;48(3):514–6.

62. Koch T, Shim I, Lindow M, Orum H, Bohr HG. Quantum mechanical studies of DNA and LNA. Nucleic Acid Ther. 2014 Apr;24(2):139–48.

63. Mann M, Wright PR, Backofen R. IntaRNA 2.0: enhanced and customizable prediction of RNA-RNA interactions. Nucleic Acids Res. 2017 Jul 3;45(W1):W435–W9.

64. Milewski MC, Kamel K, Kurzynska-Kokorniak A, Chmielewski MK, Figlerowicz M. EvOligo: A Novel Software to Design and Group Libraries of Oligonucleotides Applicable for Nucleic Acid-Based Experiments. J Comput Biol. 2017 Oct;24(10):1014–28.

65. Watanabe T, Takeda A, Mise K, Okuno T, Suzuki T, Minami N, et al. Stage-specific expression of microRNAs during Xenopus development. FEBS Lett. 2005 Jan 17;579(2):318–24.

66. Paturi S, Deshmukh MV. A Glimpse of “Dicer Biology” Through the Structural and Functional Perspective. Front Mol Biosci. 2021;8:643657.

67. Zhang H, Kolb FA, Brondani V, Billy E, Filipowicz W. Human Dicer preferentially cleaves dsRNAs at their termini without a requirement for ATP. EMBO J. 2002 Nov 1;21(21):5875–85.

68. Wallaert A, Van Loocke W, Hernandez L, Taghon T, Speleman F, Van Vlierberghe P. Comprehensive miRNA expression profiling in human T-cell acute lymphoblastic leukemia by small RNA-sequencing. Sci Rep. 2017 Aug 11;7(1):7901.

69. Kurzynska-Kokorniak A, Koralewska N, Tyczewska A, Twardowski T, Figlerowicz M. A New Short Oligonucleotide-Based Strategy for the Precursor-Specific Regulation of microRNA Processing by Dicer. PLoS One. 2013;8(10):e77703.

70. Koralewska N, Hoffmann W, Pokornowska M, Milewski M, Lipinska A, Bienkowska-Szewczyk K, et al. How short RNAs impact the human ribonuclease Dicer activity: putative regulatory feedback-loops and other RNA-mediated mechanisms controlling microRNA processing. Acta Biochim Pol. 2016;63(4):773–83.

71. Hirose G, Jacobson M. Clonal organization of the central nervous system of the frog. I. Clones stemming from individual blastomeres of the 16-cell and earlier stages. Dev Biol. 1979 Aug;71(2):191–202.

72. Falk J, Drinjakovic J, Leung KM, Dwivedy A, Regan AG, Piper M, et al. Electroporation of cDNA/Morpholinos to targeted areas of embryonic CNS in Xenopus. BMC Dev Biol. 2007 Sep 27;7:107.

73. Carrella S, D’Agostino Y, Barbato S, Huber-Reggi SP, Salierno FG, Manfredi A, et al. miR-181a/b control the assembly of visual circuitry by regulating retinal axon specification and growth. Dev Neurobiol. 2015 Nov;75(11):1252–67.

74. Campbell DS, Regan AG, Lopez JS, Tannahill D, Harris WA, Holt CE. Semaphorin 3A elicits stage-dependent collapse, turning, and branching in Xenopus retinal growth cones. J Neurosci. 2001 Nov 1;21(21):8538–47.

75. Zhao Y, Ishibashi S, Amaya E. Reverse genetic studies using antisense morpholino oligonucleotides. Methods Mol Biol. 2012;917:143–54.

76. Guo L, Lu Z. The fate of miRNA* strand through evolutionary analysis: implication for degradation as merely carrier strand or potential regulatory molecule? PLoS One. 2010 Jun 30;5(6):e11387.

77. Medley JC, Panzade G, Zinovyeva AY. microRNA strand selection: Unwinding the rules. Wiley Interdiscip Rev RNA. 2020 May;12(3):e1627.

78. Winter J, Link S, Witzigmann D, Hildenbrand C, Previti C, Diederichs S. Loop-miRs: active microRNAs generated from single-stranded loop regions. Nucleic Acids Res. 2013 May 1;41(10):5503–12.

79. Tomasello L, Distefano R, Nigita G, Croce CM. The MicroRNA Family Gets Wider: The IsomiRs Classification and Role. Front Cell Dev Biol. 2021;9:668648.

80. Langmead B, Trapnell C, Pop M, Salzberg SL. Ultrafast and memory-efficient alignment of short DNA sequences to the human genome. Genome Biol. 2009;10(3):R25.

81. Nieuwkoop PD, Faber J. Normal table of Xenopus laevis (Daudin): a systematical and chronological survey of the development from the fertilized egg till the end of metamorphosis. Garland Science; 1994.

82. Schmittgen TD, Lee EJ, Jiang J. High-throughput real-time PCR. Methods Mol Biol. 2008;429:89–98.

83. Liu Z, Wang J, Cheng H, Ke X, Sun L, Zhang QC, et al. Cryo-EM Structure of Human Dicer and Its Complexes with a Pre-miRNA Substrate. Cell. 2018 May 31;173(6):1549–50.

84. Ciechanowska K, Pokornowska M, Kurzynska-Kokorniak A. Genetic Insight into the Domain Structure and Functions of Dicer-Type Ribonucleases. Int J Mol Sci. 2021 Jan 9;22(2).

