## Supplementary Information for "Short 2′-O-methyl/LNA oligomers as highly-selective inhibitors of miRNA production *in vitro* and *in vivo*"

**This PDF file includes the following:**

Supplementary Figures S1 to S8

Legend for Supplementary Table S1

Supplementary Table S2

Supplementary Table S3

**Other supplementary materials for this manuscript include:**

Supplementary Table S1

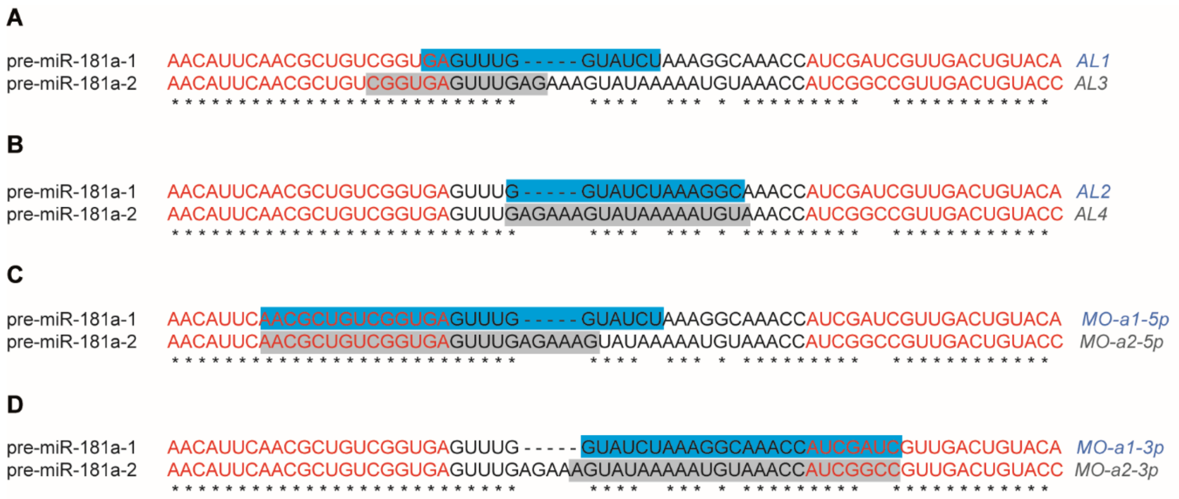


**Supplementary Figure S1. The comparison of the identity of target sequences for all inhibitors used in the study**

**(A-D)** The alignment of pre-miR-181a-1 and pre-miR-181a-2 generated in Clustal Omega. Target sites for (A) AL1 and AL3, (B) AL2 and AL4, (C) MOs-5p, (D) MOs-3p are marked with the highlights. Red font indicates the sequences of miRNAs.

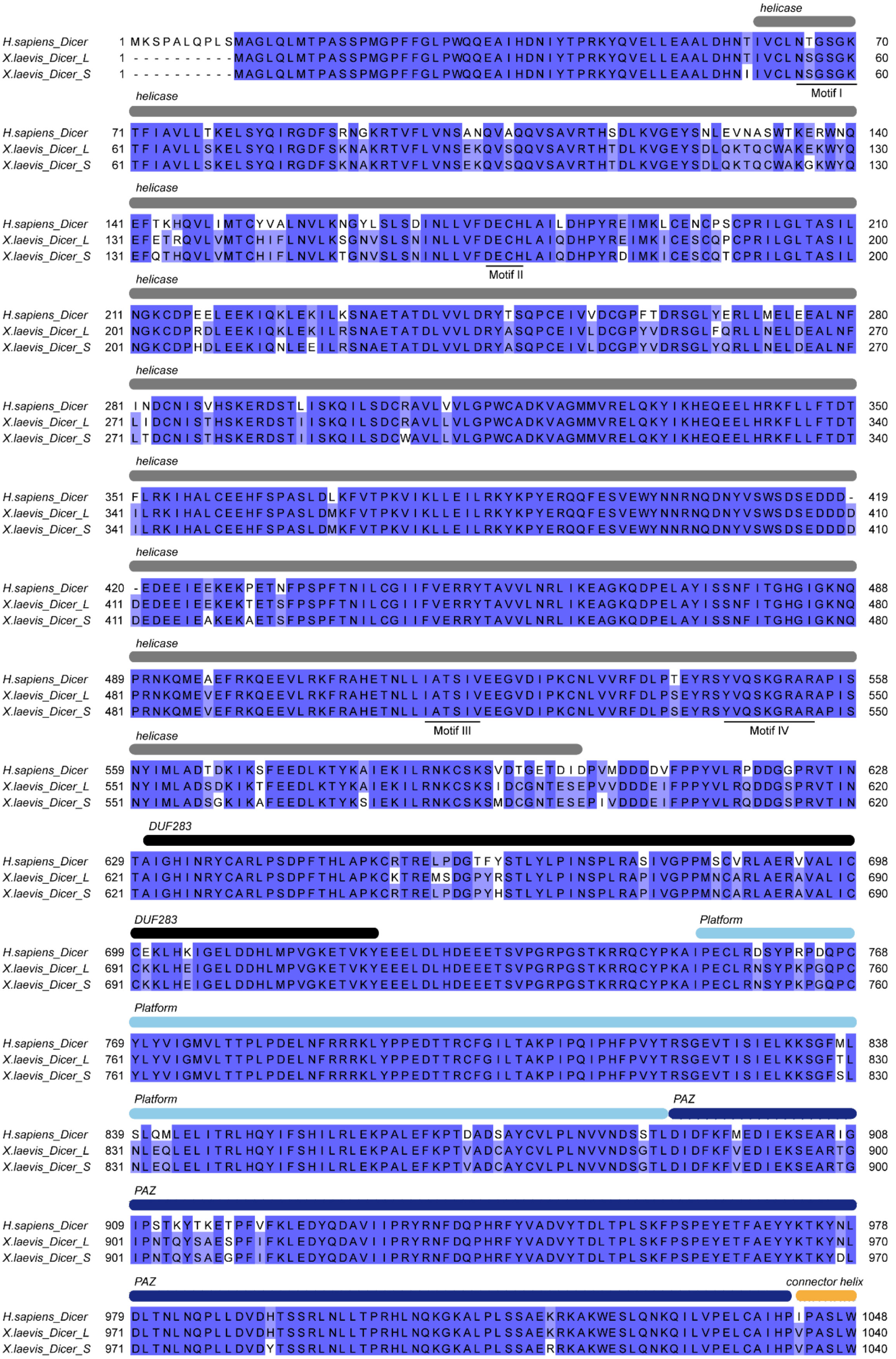
helicase
H.sapiens_Dicer 1 MKSPALQPLSMAGLQLMTPASSPMGPFFGLPWQQEAIHDNIYTPRKYQVELLEAALDHNTIVCLNTGSGK 70
X.laevis_Dicer_L 1 ----------MAGLQLMTPASSPMGPFFGLPWQQEAIHDNIYTPRKYQVELLEAALDHNTIVCLNSGSGK 60
X.laevis_Dicer_S 1 ----------MAGLQLMTPASSPMGPFFGLPWQQEAIHDNIYTPRKYQVELLEAALDHNIIVCLNSGSGK 60
Motif I
helicase
H.sapiens_Dicer 71 TFIAVLLTKELSYQIRGDFSRNGKRTVFLVNSANQVAQQVSAVRTHSDLKVGEYSNLEVNASWTKERWNQ 140
X.laevis_Dicer_L 61 TFIAVLLSKELSYQIRGDFSKNAKRTVFLVNSEKQVSQQVSAVRTHTDLKVGEYSDLQKTQCWAKEKWYQ 130
X.laevis_Dicer_S 61 TFIAVLLSKELSYQIRGDFSKNAKRTVFLVNSEKQVSQQVSAVRTHTDLKVGEYSDLQKTQCWAKGKWYQ 130
helicase
H.sapiens_Dicer 141 EFTKHQVLIMTCYVALNVLKNGYLSLSDINLLVFDECHLAILDHPYREIMKLCENCPSCPRILGLTASIL 210
X.laevis_Dicer_L 131 EFETRQVLVMTCHIFLNVLKSGNVSLSNINLLVFDECHLAIQDHPYREIMKICESCQPCPRILGLTASIL 200
X.laevis_Dicer_S 131 EFQTHQVLVMTCHIFLNVLKTGNVSLSNINLLVFDECHLAIQDHPYRDIMKICESCQTCPRILGLTASIL 200
Motif II
helicase
H.sapiens_Dicer 211 NGKCDPEELEEKIQKLEKILKSNAETATDLVVLDRYTSQPCEIVVDCGPFTDRSGLYERLLMELEEALNF 280
X.laevis_Dicer_L 201 NGKCDPRDLEEKIQKLEKILRSNAETATDLVVLDRYASQPCEIVLDCGPYVDRSGLFQRLLNELDEALNF 270
X.laevis_Dicer_S 201 NGKCDPHDLEEKIQNLEEILRSNAETATDLVVLDRYASQPCEIVLDCGPYVDRSGLYQRLLNELDEALNF 270
helicase
H.sapiens_Dicer 281 INDCNISVHSKERDSTLISKQILSDCRAVLVVLGPWCADKVAGMMVRELQKYIKHEQEELHRKFLLFTDT 350
X.laevis_Dicer_L 271 LIDCNISTHSKERDSTIISKQILSDCRAVLLVLGPWCADKVAGMMVRELQKYIKHEQEELHRKFLLFTDT 340
X.laevis_Dicer_S 271 LTDCNISTHSKERDSTIISKQILSDCWAVLLVLGPWCADKVAGMMVRELQKYIKHEQEELHRKFLLFTDT 340
helicase
H.sapiens_Dicer 351 FLRKIHALCEEHFSPASLDLKFVTPKVIKLLEILRKYKPYERQQFESVEWYNNRNQDNYVSWSDSEDDD- 419
X.laevis_Dicer_L 341 ILRKIHALCEEHFSPASLDMKFVTPKVIKLLEILRKYKPYERQQFESVEWYNNRNQDNYVSWSDSEDDDD 410
X.laevis_Dicer_S 341 ILRKIHALCEEHFSPASLDMKFVTPKVIKLLEILRKYKPYERQQFESVEWYNNRNQDNYVSWSDSEDDDD 410
helicase
H.sapiens_Dicer 420 -EDEEIEEKEKPETNFPSPFTNILCGIIFVERRYTAVVLNRLIKEAGKQDPELAYISSNFITGHGIGKNQ 488
X.laevis_Dicer_L 411 DEDEEIEEKEKTETSFPSPFTNILCGIIFVERRYTAVVLNRLIKEAGKQDPELAYISSNFITGHGIGKNQ 480
X.laevis_Dicer_S 411 DEDEEIEAKEKAETSFPSPFTNILCGIIFVERRYTAVVLNRLIKEAGKQDPELAYISSNFITGHGIGKNQ 480
helicase
H.sapiens_Dicer 489 PRNKQMEAEFRKQEEVLRKFRAHETNLLIATSIVEEGVDIPKCNLVVRFDLPTEYRSYVQSKGRARAPIS 558
X.laevis_Dicer_L 481 PRNKQMEVEFRKQEEVLRKFRAHETNLLIATSIVEEGVDIPKCNLVVRFDLPSEYRSYVQSKGRARAPIS 550
X.laevis_Dicer_S 481 PRNKQMEVEFRKQEEVLRKFRAHETNLLIATSIVEEGVDIPKCNLVVRFDLPSEYRSYVQSKGRARAPIS 550
Motif III
Motif IV
helicase
H.sapiens_Dicer 559 NYIMLADTDKIKSFEEDLKTYKAIEKILRNKCSKSVDTGETDIDPVMDDDDVFPPYVLRPDDGGPRVTIN 628
X.laevis_Dicer_L 551 NYIMLADSDKIKTFEEDLKTYKAIEKILRNKCSKSIDCGNTESEPVVDDDEIFPPYVLRQDDGSPRVTIN 620
X.laevis_Dicer_S 551 NYIMLADSGKIKAFEEDLKTYKSIEKILRNKCSKSMDCGNTESEPIVDDDEIFPPYVLRQDDGSPRVTIN 620
DUF283
H.sapiens_Dicer 629 TAIGHINRYCARLPSDPFTHLAPKCRTRELPDGTFYSTLYLPINSPLRASIVGPPMSCVRLAERVVALIC 698
X.laevis_Dicer_L 621 TAIGHINRYCARLPSDPFTHLAPKCKTREMSDGPYRSTLYLPINSPLRAPIVGPPMNCARLAERAVALIC 690
X.laevis_Dicer_S 621 TAIGHINRYCARLPSDPFTHLAPKCRTRELPDGPYHSTLYLPINSPLRAPIVGPPMNCARLAERAVALIC 690
DUF283
Platform
H.sapiens_Dicer 699 CEKLHKIGELDDHLMPVGKETVKYEEELDLHDEEETSVPGRPGSTKRRQCYPKAIPECLRDSYPRPDQPC 768
X.laevis_Dicer_L 691 CKKLHEIGELDDHLMPVGKETVKYEEELDLHDEEETSVPGRPGSTKRRQCYPKAIPECLRNSYPKPGQPC 760
X.laevis_Dicer_S 691 CKKLHEIGELDDHLMPVGKETVKYEEELDLHDEEETSVPGRPGSTKRRQCYPKAIPECLRNSYPKPGQPC 760
Platform
H.sapiens_Dicer 769 YLYVIGMVLTTPLPDELNFRRRKLYPPEDTTRCFGILTAKPIPQIPHFPVYTRSGEVTISIELKKSGFML 838
X.laevis_Dicer_L 761 YLYVIGMVLTTPLPDELNFRRRKLYPPEDTTRCFGILTAKPIPQIPHFPVYTRSGEVTISIELKKSGFTL 830
X.laevis_Dicer_S 761 YLYVIGMVLTTPLPDELNFRRRKLYPPEDTTRCFGILTAKPIPQIPHFPVYTRSGEVTISIELKKSGFSL 830
Platform
PAZ
H.sapiens_Dicer 839 SLQMLELITRLHQYIFSHILRLEKPALEFKPTDADSAYCVLPLNVVNDSSTLDIDFKFMEDIEKSEARIG 908
X.laevis_Dicer_L 831 NLEQLELITRLHQYIFSHILRLEKPALEFKPTVADCAYCVLPLNVVNDSGTLDIDFKFVEDIEKSEARTG 900
X.laevis_Dicer_S 831 NLEQLELITRLHQYIFSHILRLEKPALEFKPTVADCAYCVLPLNVVNDSGTLDIDFKFVEDIEKSEARTG 900
PAZ
H.sapiens_Dicer 909 IPSTKYTKETPFVFKLEDYQDAVIIPRYRNFDQPHRFYVADVYTDLTPLSKFPSPEYETFAEYYKTKYNL 978
X.laevis_Dicer_L 901 IPNTQYSAESPFIFKLEDYQDAVIIPRYRNFDQPHRFYVADVYTDLTPLSKFPSPEYETFAEYYKTKYNL 970
X.laevis_Dicer_S 901 IPNTQYSAEGPFIFKLEDYQDAVIIPRYRNFDQPHRFYVADVYTDLTPLSKFPSPEYETFAEYYKTKYDL 970
PAZ
connector helix
H.sapiens_Dicer 979 DLTNLNQPLLDVDHTSSRLNLLTPRHLNQKGKALPLSSAEKRKAKWESLQNKQILVPELCAIHPIPASLW 1048
X.laevis_Dicer_L 971 DLTNLNQPLLDVDHTSSRLNLLTPRHLNQKGKALPLSSAEKRKAKWESLQNKQILVPELCAIHPVPASLW 1040
X.laevis_Dicer_S 971 DLTNLNQPLLDVDYTSSRLNLLTPRHLNQKGKALPLSSAERRKAKWESLQNKQILVPELCAIHPVPASLW 1040

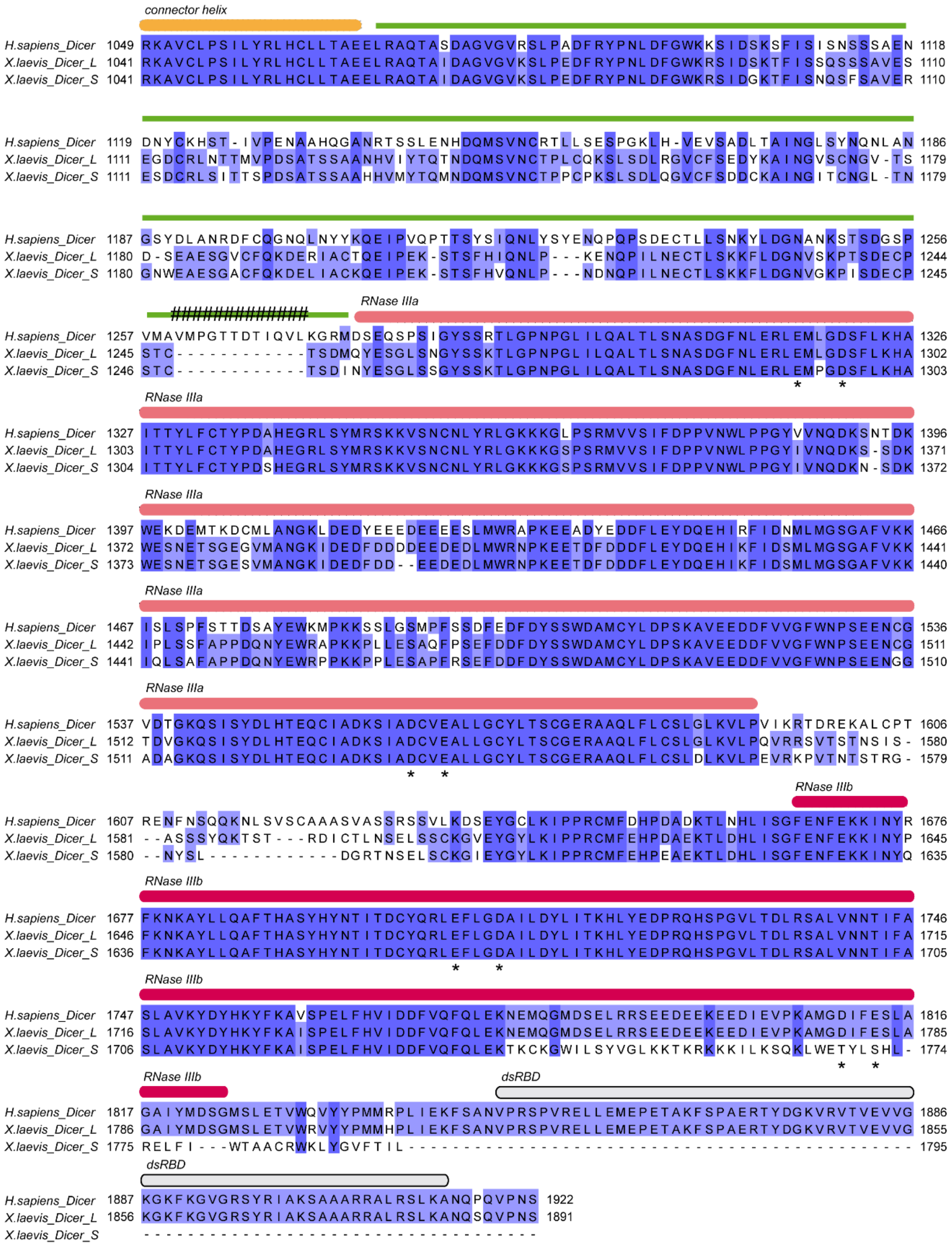

**Supplementary Figure S2. The comparison of the sequences of human and *X. laevis* Dicer proteins**

Multiple sequence alignment performed using ClustalW [1]. Sequences retrieved from UniProtKB: Q9UPY3, A0A1L8F9L1, D0UED5. Graded blue highlights of the sequence indicate the level of amino acid conservation. Protein domains are marked above the sequences. Colour code as in Figure 2. Asterisks (*) indicate amino acids of the catalytic core; hashtags (#) point a 12-aa difference within the molecular ruler helix among human and *X. laevis* proteins. Abbreviations: DUF283, domain of unknown function 283; PAZ, Piwi/Argonaute/Zwille domain; dsRBD, dsRNA binding domain.

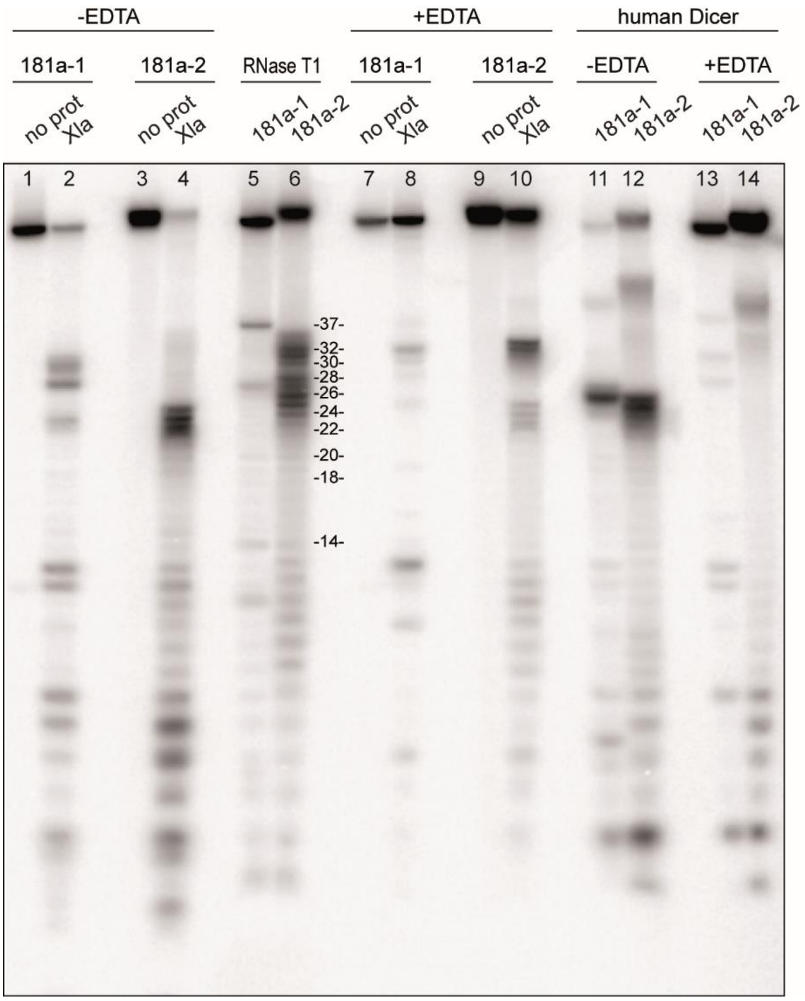


**Supplementary Figure S3. Characterization of the pre-miRNA cleavage pattern generated in cytosolic extracts**

Uncropped image of a representative gels presented in Figure 2, showing the full range of products generated upon the incubation of 5′-$^{32}$P-labeled pre-miRNA with *Xenopus* cytosolic extracts (lines 1-4, 7-10) or human Dicer (lines 11-14), with or without EDTA. Partial hydrolysis of RNA by RNase T1 was used to generate a ladder (lines 5, 6). Reproducible results were obtained using different batches of the extracts.

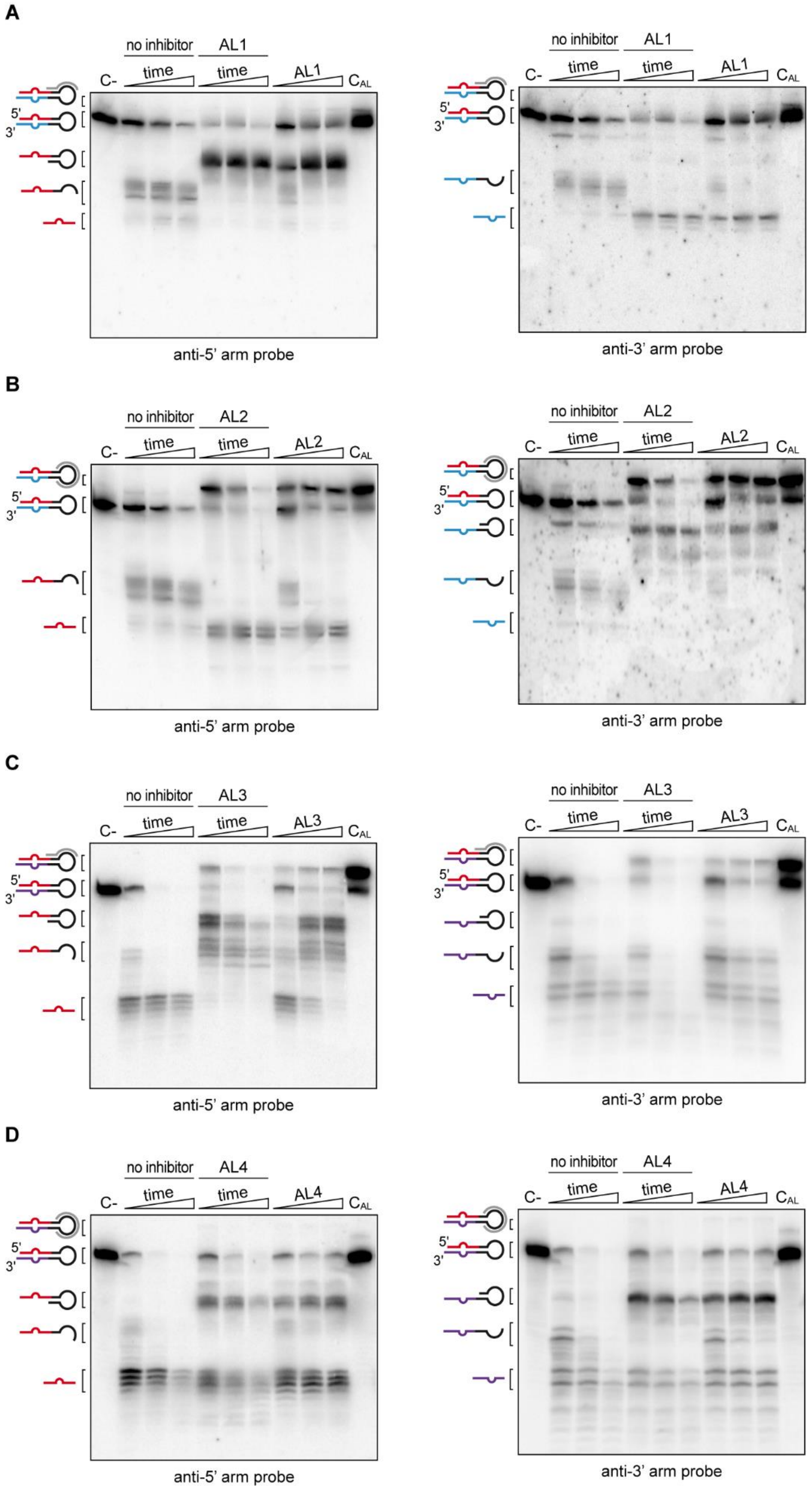
A
no inhibitor
AL1
C-
time
time
AL1
CAL
5'
3'
anti-5' arm probe
anti-3' arm probe
B
no inhibitor
AL2
C-
time
time
AL2
CAL
5'
3'
anti-5' arm probe
anti-3' arm probe
C
no inhibitor
AL3
C-
time
time
AL3
CAL
5'
3'
anti-5' arm probe
anti-3' arm probe
D
no inhibitor
AL4
C-
time
time
AL4
CAL
5'
3'
anti-5' arm probe
anti-3' arm probe

**Supplementary Figure S4. Validation of pre-miR-181a-1 and pre-miR-181a-2 cleavage pattern in the presence of 2′-OMe/LNA ASOs targeting 5′ arm of the precursors**

**(A-D)** Northern blot analysis of the cleavage pattern of pre-miR-181a-1 in the presence of AL1 (A) or AL2 (B), and pre-miR-181a-2 in the presence of AL3 (C) or AL4 (D). RNA was incubated with 100 molar excess of the indicated AL and *Xenopus* cytosolic extract for 0.5, 1.5, 3 h (time change indicated by a triangle) or with 1, 10 or 100 molar excess of the given AL for 0.5 h (AL concentration change indicated by a triangle). RNA was visualized using DNA probes targeting either 5′ or 3′ arm of the pre-miRNA as indicated. Schematic representation of the pre-miRNA, AL and cleavage products is given on the left to the blots. Abbreviations: C-, pre-miRNA incubated in a buffer with no AL nor protein; $C_{AL}$, pre-miRNA incubated in a buffer with AL but without protein.

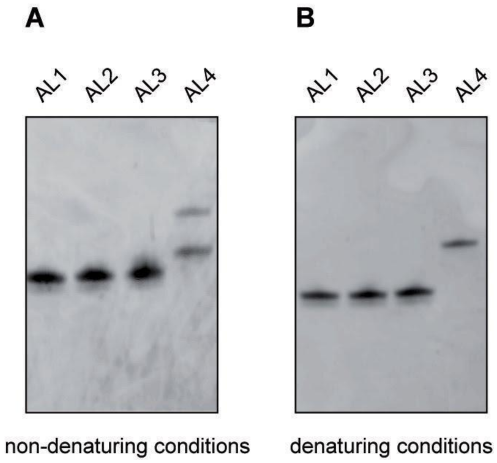


**Supplementary Figure S5. Assessment of RNA secondary structure forms adopted by designed 2′-OMe/LNA oligomers**

**(A)** RNA folding analyzed by non-denaturing PAGE of the unlabeled 2′-OMe/LNA antisense oligomers (AL1-4), followed by SYBR Gold staining. **(B)** RNA integrity verified by denaturing PAGE of the unlabeled 2′-OMe/LNA ASOs (AL1-4), followed by SYBR Gold staining.

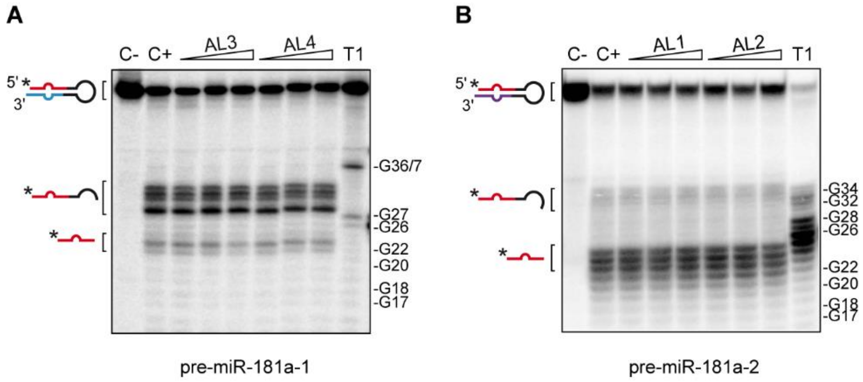


**Supplementary Figure S6.** ***In vitro*** **validation of the target-specificity of the designed 2′-OMe/LNA ASOs**

**(A, B)** 5′-$^{32}$P-labeled pre-miR-181a-1 (A) and pre-miR-181a-2 (B) were incubated with hDicer in the presence of AL2 or AL3, and AL1 or AL2, as indicated. Triangles represent increasing amounts of the indicated AL (pre-miRNA:oligomer molar ratios of 1:1, 1:10, and 1:100). Abbreviations: C-, sample with no protein, nor inhibitor added; C+, sample with human Dicer, without any inhibitor; T1, RNase T1 ladder.

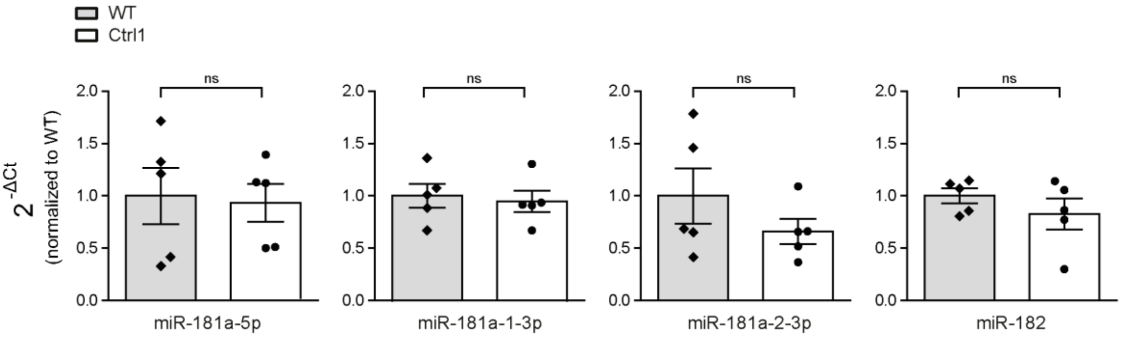


**Supplementary Figure S7.** ***In vivo*** **evaluation of 2′-OMe/LNA control oligomer**
miRNAs expression levels quantified using the $2^{-\Delta Ct}$ method and U6 as normalizer, after Ctrl oligomer microinjection. Data are normalized to WT control embryos. Data information: Values are mean ± SEM. Statistics: n=5 independent experiments, each data point represents a single RT–qPCR, unpaired t-test. Abbreviations: WT, wild type; ns, not significant.

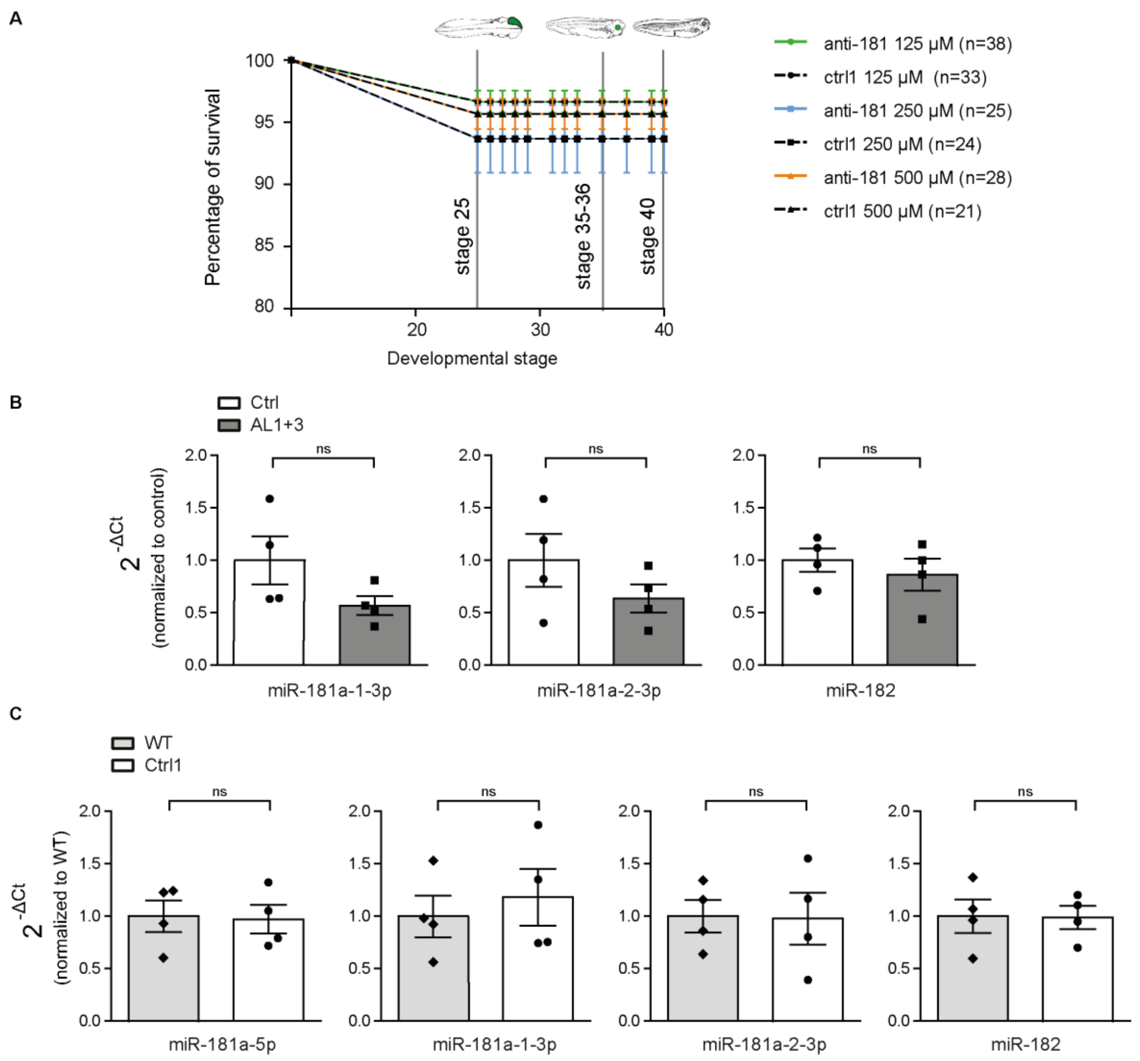


**Supplementary Figure S8.** ***In vivo*** **evaluation of 2´-OMe/LNA ASOs compatibility**
**(A)** Survival embryos rate after inhibitors eye delivery, in controls and AL1+3. **(B, C)** miRNAs expression levels quantified using the $2^{-\Delta Ct}$ method and U6 as normalizer. Data are normalized to control (B) or to WT (C). Data information: Values are mean ± SEM. Statistics: n=3 independent experiments for each of the concentrations tested (A), total number of embryos is reported in parenthesis (A); n=5 independent experiments, each data point represents a single RT–qPCR, unpaired t-test (B, C). Abbreviations: ns, not significant; WT, wild type.

**Supplementary Table S1.** Characterization of the interaction between the oligomers designed in the study and potential targets found within *X. laevis* miRNA and pre-miRNA (a separate Excel file).

**Supplementary Table S2.** Sequences (5′-3′) of the best 162 negative control oligomers that can be used in *X. laevis* model. The oligomer used in the study is marked in red.

| | | | |
|---|---|---|---|
| 1 CGUAUACUUCGCG | 53 CUACCGCGUAAGG | 105 GUUAGGUCGCGUA | 157 UCGCGUACCGAGU |
| 2 UCGCGCCACGAUA | 54 CCUUACGCGGUAG | 106 CGUCGAUACUAGG | 158 ACUCGGUACGCGA |
| 3 UAUCGUGGCGCGA | 55 ACGCCUAGCGUAA | 107 CCUAGUAUCGACG | 159 UUACGCGCCGUAA |
| 4 CUACGCGGCGUAA | 56 UACGCGAAUCGUU | 108 UACGCGAUCCGAU | 160 UUACGGCGCGUAA |
| 5 UACGCGGCGUAAC | 57 AACGAUUCGCGUA | 109 AUCGGAUCGCGUA | 161 UCGCGGAGUACUA |
| 6 GUUACGCCGCGUA | 58 UAAUCGCGUACCG | 110 CGCGACCUAUCGU | 162 UAGUACUCCGCGA |
| 7 CUCGCGAAUAACG | 59 CGGUACGCGAUUA | 111 ACGAUAGGUCGCG | |
| 8 UCGCGUAUUCGCG | 60 UUAGCGCGGUACG | 112 GUUCGUAACGCGC | |
| 9 CGCGAAUACGCGA | 61 CGUACCGCGCUAA | 113 GCGCGUUACGAAC | |
| 10 UACGCGCAUCGUA | 62 UUCGACGCGAGUA | 114 CGUUCGCGUAGGU | |
| 11 UAUUACGCGAGCG | 63 UACUCGCGUCGAA | 115 ACCUACGCGAACG | |
| 12 CGCUCGCGUAAUA | 64 UAAGUCGACGCGU | 116 CUACGUACGCUCG | |
| 13 UUACGUUACGCGA | 65 UACGACGGUAAGC | 117 UCGGCGUAGUACC | |
| 14 UCGCGUAACGUAA | 66 ACGCGUCGACUUA | 118 GGUACUACGCCGA | |
| 15 CUAUCGCGCUAGU | 67 GCUUACCGUCGUA | 119 GUUACGAUCGCGC | |
| 16 CGCGAAUUAAUCG | 68 UCGUACGCGAUUG | 120 UUCGUACCGCUAG | |
| 17 CGUUAGUACGCCG | 69 CGUCGUUAGGACG | 121 CUAGCGGUACGAA | |
| 18 CGGCGUACUAACG | 70 CGUCCUAACGACG | 122 UCGUUAGGUCGCG | |
| 19 ACUAGCGCGAUAG | 71 UAUCGACGCGUAG | 123 CGGUCGUAUACCG | |
| 20 UAAGCGCGUAACG | 72 CUACGCGUCGAUA | 124 CGGUAUACGACCG | |
| 21 CGUUACGCGCUUA | 73 CAAUCGCGUACGA | 125 CGCGACCUAACGA | |
| 22 CGUAACGAACCGU | 74 CGUCGUCGCGAAU | 126 GCACGCUAGUACG | |
| 23 CCGCGUAGCGUAU | 75 AUUCGCGACGACG | 127 CGUACUAGCGUGC | |
| 24 AUACGCUACGCGG | 76 GCUAGUCUACGCG | 128 CGAUACCGUACGC | |
| 25 ACGGUUCGUUACG | 77 CGCGUAGACUAGC | 129 GGACGUACGCGAU | |
| 26 UUACGCGACGCGU | 78 CGCGAUACGAUCU | 130 CGACUAACGGUCG | |
| 27 CUAUCUACGCUCG | 79 AGAUCGUAUCGCG | 131 CGACCGUUAGUCG | |
| 28 CGAGCGUAGAUAG | 80 GUCGCGUACCGUU | 132 CGUCGAUACGUCG | |
| 29 ACGCGUCGCGUAA | 81 CGCCGUACGUAAU | 133 CGACGUAUCGACG | |
| 30 UUACGCGAACCUA | 82 AUUACGUACGGCG | 134 CGUUCGCGUAUCG | |
| 31 UAGGUUCGCGUAA | 83 AACGGUACGCGAC | 135 CGAUACGCGAACG | |
| 32 UUACGCCGCGUAG | 84 UCGCGCAAUCUUA | 136 CUAAUUCGCGACG | |
| 33 CGUUCGCGUAAGU | 85 GUUCGCGCCUAAG | 137 CGUCGCGAAUUAG | |
| 34 CUACCGAUUAGCG | 86 CUUAGGCGCGAAC | 138 GCGUACUACGACG | |
| 35 ACUUACGCGAACG | 87 UUAGUCGAGCGCG | 139 CGUCGUAGUACGC | |
| 36 CGCUAAUCGGUAG | 88 UACGGCGCGUAAC | 140 UCACGAUUGCGCG | |
| 37 UAACUAGCUCGCG | 89 GUUACGCGCCGUA | 141 CGCGCAAUCGUGA | |
| 38 UAACUCGCGGACG | 90 CGCGCUCGACUAA | 142 CGCGGUCGAUAGU | |
| 39 GUACGCGAGUACG | 91 UAAGUUCGCGACG | 143 ACUAUCGACCGCG | |
| 40 CGUACUCGCGUAC | 92 CGUCGCGAACUUA | 144 GUCGAUCGACGAU | |
| 41 CGUCCGCGAGUUA | 93 UACGCGAAGUCCG | 145 GUCGACAUCGCGU | |
| 42 CGCGUACUACCGU | 94 CGGACUUCGCGUA | 146 UCGACUAACGUCG | |
| 43 ACGGUAGUACGCG | 95 UACUAGCGUACCG | 147 CGACGUUAGUCGA | |
| 44 CCGCGACUUAGUU | 96 CGGUACGCUAGUA | 148 UAUAUCGACGCGU | |
| 45 AACUAAGUCGCGG | 97 UACGCGACUAGCC | 149 ACGCGUCGAUAUA | |
| 46 UCGACCGCGUAAG | 98 UGCGCGUCGAAUG | 150 GUUACGCGAACCG | |
| 47 CUUACGCGGUCGA | 99 UUAACGGUACGCG | 151 CGGUUCGCGUAAC | |
| 48 UACGCGUCGAUAG | 100 CAUUCGACGCGCA | 152 ACUAGGUACGCGU | |
| 49 CUAUCGACGCGUA | 101 CGCGUACCGUUAA | 153 ACGCGUACCUAGU | |
| 50 CGGUUCGACGUAU | 102 GUACGCGUUAGUU | 154 UACUCGCGACCAC | |
| 51 AUACGUCGAACCG | 103 AACUAACGCGUAC | 155 UAAGAUCGCGUAG | |
| 52 UUACGCUAGGCGU | 104 UACGCGACCUAAC | 156 CUACGCGAUCUUA | |

### Supplementary Table S3. Oligonucleotides used in the study

| **Oligonucleotide name** | **Sequence (5′→ 3′) or Catalog and assay number** |
|---|---|
| 2′-O-methyl/LNA (in-house) | |
| AL1 | AGAUACCAAACUC |
| AL2 | GCCUUUAGAUACC |
| AL3 | CUCAAACUCACCG |
| AL4 | ACAUUUUUAUACUUUCUC |
| Ctrl | CGUAUACUUCGCG |
| DNA (IBB PAS) | |
| pre-miR-181a anti-5p arm probe | ACTCACCGACAGCGTTGAATGTT |
| pre-miR-181a anti-3p arm probe | GGTACAGTCAACGGCCGATGGT |
| RNA (FUTUREsynthesis) | |
| pre-miR-181a-1 | AACAUUCAACGCUGUCGGUGAGUUUGGUAUCUAAAGGC AAACCAUCGAUCGUUGACUGUACA |
| pre-miR-181a-2 | AACAUUCAACGCUGUCGGUGAGUUUGAGAAAGUAUAAA AAUGUAAACCAUCGGCCGUUGACUGUACC |
| Morpholino (GeneTools) | |
| MO-a1-5p | AGATACCAAACTCACCGACAGCGTT |
| MO-a2-5p | CTTTCTCAAACTCACCGACAGCGTT |
| MO-a1-3p | GATCGATGGTTTGCCTTTAGATAC |
| MO-a2-3p | GGCCGATGGTTTATATTTTTATACT |
| TaqMan MicroRNA Assay (ThermoFisher) | |
| miR-181a-5p | Cat# 4427975; 000480 |
| miR-181a-1-3p | Cat# 4440886; 004367 |
| miR-181a-2-3p | Cat# 4440886; 005555 |
| miR-182 | Cat# 4427975; 000597 |
| snU6 | Cat# 4427975; 001973 |

References

1. Thompson JD, Gibson TJ, Higgins DG. Multiple sequence alignment using ClustalW and ClustalX. Curr Protoc Bioinformatics. 2002 Aug;Chapter 2:Unit 2 3.
2. Liu Z, Wang J, Cheng H, Ke X, Sun L, Zhang QC, et al. Cryo-EM Structure of Human Dicer and Its Complexes with a Pre-miRNA Substrate. Cell. 2018 May 17;173(5):1191-203 e12.
3. Ciechanowska K, Pokornowska M, Kurzynska-Kokorniak A. Genetic Insight into the Domain Structure and Functions of Dicer-Type Ribonucleases. Int J Mol Sci. 2021 Jan 9;22(2).
